# Deep multiple-instance learning accurately predicts gene haploinsufficiency and deletion pathogenicity

**DOI:** 10.1101/2023.08.29.555384

**Authors:** Zhihan Liu, Yi-Fei Huang

## Abstract

Copy number losses (deletions) are a major contributor to the etiology of severe genetic disorders. Although haploinsufficient genes play a critical role in deletion pathogenicity, current methods for deletion pathogenicity prediction fail to integrate multiple lines of evidence for haploinsufficiency at the gene level, limiting their power to pinpoint deleterious deletions associated with genetic disorders. Here we introduce DosaCNV, a deep multiple-instance learning framework that, for the first time, models deletion pathogenicity jointly with gene haploinsufficiency. By integrating over 30 gene-level features potentially predictive of haploinsufficiency, DosaCNV shows unmatched performance in prioritizing pathogenic deletions associated with a broad spectrum of genetic disorders. Furthermore, DosaCNV outperforms existing methods in predicting gene haploinsufficiency even though it is not trained on known haploinsufficient genes. Finally, DosaCNV leverages a state-of-the-art technique to quantify the contributions of individual gene-level features to haploinsufficiency, allowing for human-understandable explanations of model predictions. Altogether, DosaCNV is a powerful computational tool for both fundamental and translational research.

## Background

Copy number variations (CNVs) are a major contributor to human genetic diversity^1,2^. These structural variations, which involve deletions or duplications of genomic regions ranging from 50 base pairs to several megabase pairs, result from diverse mutational mechanisms during DNA replication, recombination, and repair^3,4^. CNVs may provide adaptive advantages under certain circumstances^5,6^; however, they can also lead to diseases when critical genomic regions are affected^7^. Both common and rare CNVs (population frequency < 1%) contribute to disease development, with rare CNVs particularly implicated in rare genetic disorders^2,8–13^. In the context of disease-associated CNVs, pathogenicity often arises from alterations affecting dosage-sensitive (DS) driver genes^14,15^. This concept of dosage sensitivity refers to the emergence of deleterious phenotypes when gene copy number changes. Specifically, haploinsufficient (HI) genes are intolerant to copy number loss, whereas triplosensitive (TS) genes are intolerant to copy number gain^16,17^. Thus, identifying DS genes is essential for prioritizing clinically relevant CNVs and understanding the genetic mechanisms of associated diseases.

Efforts to discover DS genes primarily involve three frameworks: machine learning-based methods, case-control data-based statistical methods, and selection pressure-based approaches. Machine learning-based methods utilize gene-level features and various algorithms to differentiate manually curated DS genes from dosage-insensitive genes^18–23^. While these models identify predictive attributes of DS genes, the limited availability of high confidence training data restricts their ability to capture the full spectrum of DS genes. Alternatively, statistical methods can be utilized to analyze case-control CNV data to identify DS genes, assuming that these genes have a higher prevalence among affected individuals^15^. For instance, a recent approach employs a modified GWAS fine-mapping framework to estimate gene-level dosage sensitivity^24^. Although this framework can predict DS genes without labeled training data, it is challenging to pinpoint causal genes because CNVs often span multiple genes. Finally, selection pressure-based methods utilize natural selection against loss-of-function (LOF) mutations or CNVs as indicators of dosage sensitivity^25–30^. Although dosage sensitivity is strongly correlated with selection pressure at the gene level, exceptions do exist. For instance, alterations in pleiotropic genes critical for fitness may not lead to significant phenotypic changes^31,32^, and DS genes associated with late-onset disorders may not be subject to strong selection pressure^33–36^. Additionally, these selection pressure-based methods may be underpowered for detecting short DS genes due to sparse polymorphisms in the human genome^27,28^.

Previous research prioritizes pathogenic CNVs using two frameworks: rule-based methods and supervised machine learning methods. Rule-based approaches automate pathogenicity scoring systems based on clinical CNV interpretation guidelines^37,38^, such as the one proposed by the American College of Medical Genetics and Genomics (ACMG) and ClinGen^39^. However, these methods face limitations due to rigid score assignments and difficulties in identifying pathogenicity originating from uncatalogued genomic regions. On the other hand, supervised machine learning methods, often trained on clinically curated CNVs, utilize simple summary statistics aggregated across affected genomic elements, such as the mean score of dosage sensitivity across genes, for pathogenicity prediction. However, the optimal summary statistics for deletion pathogenicity prediction are often unknown, limiting the accuracy of supervised machine learning methods. For instance, the mean score of dosage sensitivity across genes might not adequately convey the significance of a pathogenic CNV harboring a single highly penetrant DS driver gene amidst multiple dosage-insensitive genes.

Furthermore, while current methods offer valuable insights into gene dosage sensitivity and CNV pathogenicity, they struggle to connect the two. DS gene identification methods predict gene dosage sensitivity but lack a natural way of aggregating these predictions to infer CNV pathogenicity. Conversely, pathogenic CNV classifiers predict CNV pathogenicity without identifying DS driver genes and evaluating their contributions. To bridge this gap and enhance predictions at both the gene and CNV levels, we propose that multiple-instance learning (MIL) presents a promising, unified solution for the joint predictions of gene dosage sensitivity and CNV pathogenicity^40^. MIL is suited for problems where each sample, or bag, in the dataset contains a varying number of observations, or instances, and is classified as positive if at least one instance is positive^40^. The objective is to predict labels at both the bag and instance levels, assuming that the bag-level label results from the combined effect of individual instances, even when only bag-level labels are available during training. This approach is well-aligned with our problem, as we can view each CNV as a bag and the affected genes within it as instances, with pathogenicity labels given solely at the bag (CNV) level.

In this study, we introduce DosaCNV, a deep MIL framework modeling deletion pathogenicity through the joint effect of haploinsufficiency from affected genes. We focus exclusively on deletions that affect protein-coding genes, which have more direct functional consequences compared to deletions that only affect non-coding regions and duplications. DosaCNV leverages over 30 gene-level features from diverse biological processes and is trained on clinically curated deletions. It outperforms existing methods in predicting pathogenic deletions and HI genes across various sources. DosaCNV demonstrates the potential to leverage large-scale deletion data for enhancing the predictions of HI genes and associating these predictions with deletion pathogenicity, establishing it as an accurate and valuable computational tool for both fundamental and translational research.

## Results

### Overview of DosaCNV

DosaCNV is a supervised deep MIL model designed to simultaneously infer the pathogenicity of coding deletions and the haploinsufficiency of genes, based on the assumption that the joint effect of gene haploinsufficiency determines deletion pathogenicity. Utilizing a fully connected neural network, chosen for its strong ability to automatically extract useful features, DosaCNV analyzes coding deletions by first computing the haploinsufficiency of affected genes (Fig. 1). This fully connected neural network processes an input matrix for each deletion, with rows representing affected genes and columns representing gene-level features. Then, it generates a vector of predicted HI probabilities (P_HI_) for each affected gene within the deletion (Fig. 1).

**Figure 1.**
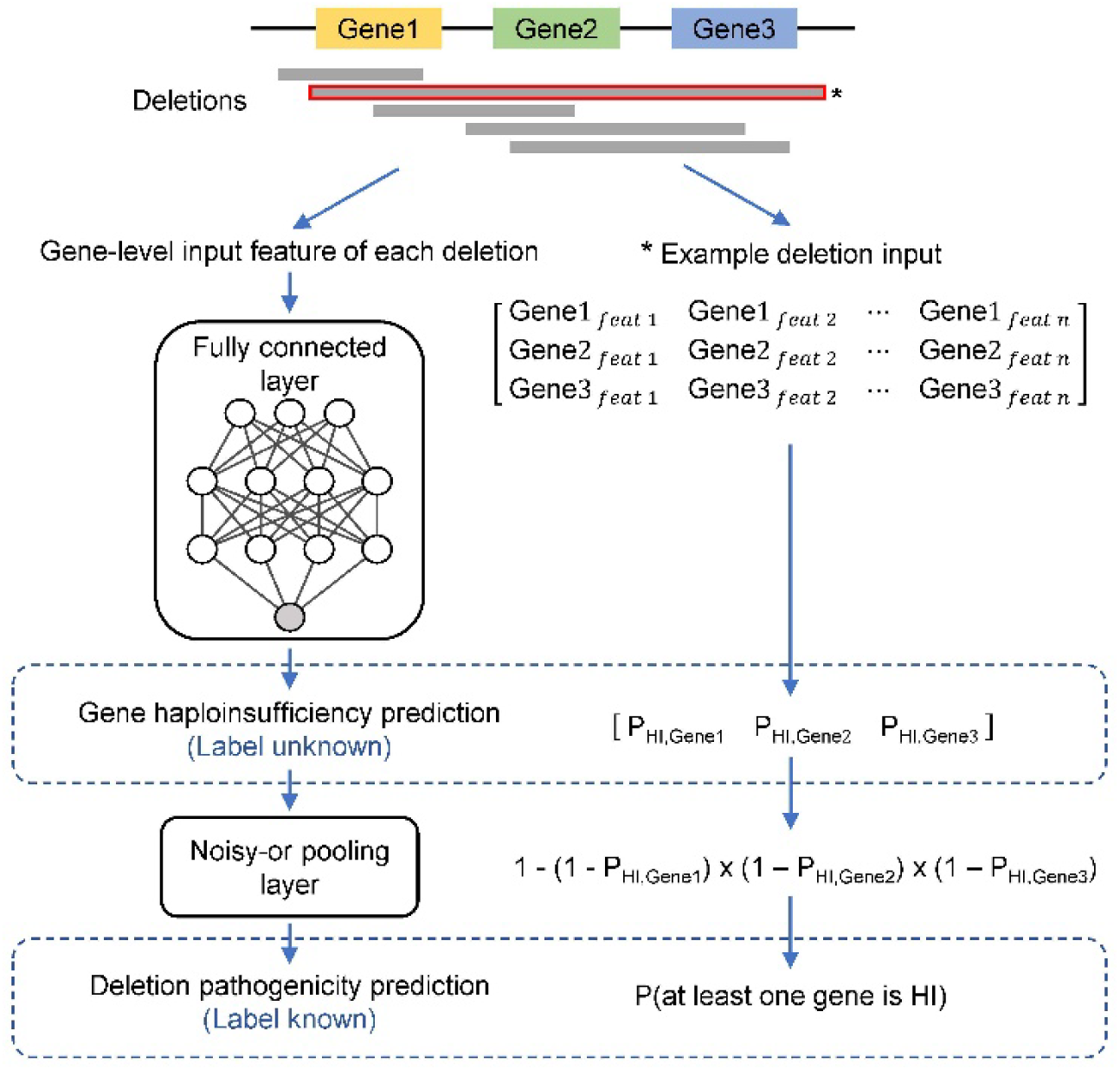
Schematic representation of the DosaCNV model. DosaCNV utilizes a fully connected architecture in conjunction with a noisy-or pooling layer to jointly infer gene haploinsufficiency and coding deletion pathogenicity. Deletions are first annotated with gene-level features of affected genes as input. For a given deletion, the fully connected architecture processes its gene-level input features and predicts the HI probability (PHI) for each affected gene. Subsequently, the noisy-or pooling layer integrates these gene-level predictions, assuming independent gene effects, to compute the probability of the deletion harboring at least one HI gene, which serves as a proxy for deletion-level pathogenicity. The right side of the panel illustrates an example deletion input, along with its intermediate gene-level predictions and the noisy-or pooling calculation.

Subsequently, DosaCNV utilizes the noisy-or pooling function^41^, a probabilistic variant of the logical-or operator commonly used in Bayesian networks, to combine the predicted HI probabilities at the gene level into a scalar output ranging from 0 to 1, representing the joint effect of gene-level haploinsufficiency (Fig. 1). This output is also the predicted probability that the given deletion overlaps with at least one HI gene by the formalization of the noisy-or function^41^, and serves as a proxy for deletion pathogenicity (Fig. 1). Notably, due to the noisy-or pooling approach, DosaCNV assumes independent effects between affected genes and does not account for gene-gene interactions.

DosaCNV’s design addresses the challenge of limited access to curated HI genes faced by previous supervised gene classifiers. By training the model’s fully connected architecture for HI prediction on all affected genes within the deletion data, DosaCNV potentially accesses a larger and more diverse agglomeration of HI genes compared to currently available sets. This approach may enable DosaCNV to capture distinct HI gene characteristics more effectively, enhancing HI gene prediction relative to other DS gene classifiers. Combining this improvement with the probabilistic integration of HI predictions for all affected genes within a deletion of interest, DosaCNV could potentially yield more accurate deletion-level pathogenicity predictions.

### Training DosaCNV on clinically curated deletions

We retrieved 6,180 pathogenic and 5,822 benign deletions overlapping with gene coding sequences (CDSs) from ClinVar^42^, a pan-disorder repository of clinically curated genetic variants. Also, we collected 33 gene-level features potentially predictive of haploinsufficiency. These gene-level features can be grouped into six categories: genomics, epigenomics, evolutionary constraints, functional genomics, deleteriousness upon perturbation, and biological networks (Supplementary Data 1). We used the features of affected protein-coding genes within a deletion as the input features of DosaCNV, and used the clinically curated pathogenicity of the deletion in ClinVar as the label. Then, we utilized a “leave-one-chromosome-out” approach to partition the data into a training set, a validation set, and a held-out test set (Methods). We trained DosaCNV on the training set and optimized the hyperparameters of DosaCNV on the validation set. To minimize potential bias arising from relying on deletion length for predictions^43,44^, we matched pathogenic and benign deletions based on the number of affected protein-coding genes within the training and validation sets. The test set remained unmatched for subsequent analysis. As a result, the final training, validation, and test sets contained 2,308, 1,060, and 2,832 coding deletions, respectively.

### Evaluating DosaCNV and alternative methods on ClinVar pathogenic deletions

We assessed DosaCNV’s performance in predicting pathogenic deletions on the held-out test set. To mitigate potential biases in performance assessment due to class imbalance and length differences between pathogenic and benign deletions, we benchmarked DosaCNV with five alternative methods (TADA^44^, StrVCTVRE^45^, HIS-LOD^18^, X-CNV^46^, and CADD-SV^47^) on the unmatched held-out test set and three subsets matched for the number of affected protein-coding genes, deletion length, and both criteria, respectively.

Within the unmatched test set, 1,211 pathogenic deletions and 1,280 benign deletions had predictions from all five alternative methods (n = 2,491). DosaCNV achieved the highest area under the receiver-operating-characteristic curve (AUC) of 0.968 (Fig. 2A; Supplementary Table 1). To evaluate the performance of various methods for deletions with different lengths, we categorized the unmatched test set into four length groups: small (<50 kb), medium (50-100 kb), large (100-1000 kb), and extra-large (>1000 kb). DosaCNV consistently delivered superior or comparable predictions across all length groups (Fig. 2B; Supplementary Table 2). Although other methods also performed reasonably well, their performance might be attributed to differences in length distribution between pathogenic and benign deletions (Supplementary Fig. 1A & B). Indeed, using either the number of affected genes or deletion length alone as a predictor for deletion pathogenicity yielded AUC values as high as 0.886 and 0.912, respectively (Fig. 2A).

**Figure 2.**
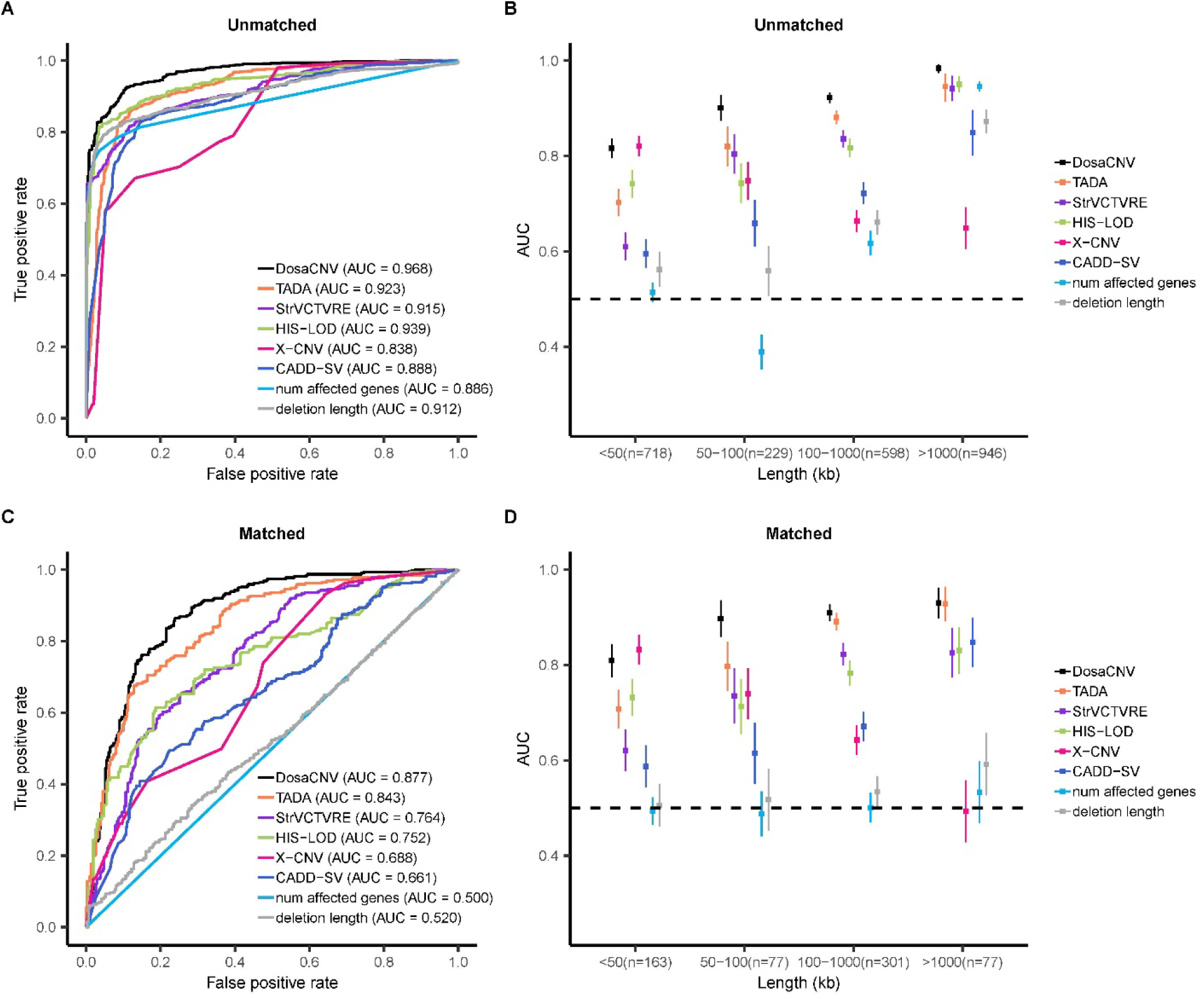
Performance comparison of DosaCNV and alternative methods in predicting pathogenicity of deletions from the unmatched and matched ClinVar held-out test sets. For each set, comparisons are presented for the entire set and four deletion subgroups based on length: small (<50 kb), medium (50-100 kb), large (100-1000 kb), and extra-large (>1000 kb). (A)(B) Unmatched test set (n=2,491). (C)(D) Test set matched by both the number of affected genes and deletion length (n=618). Vertical lines denote ± one standard error. Horizontal dashed lines indicate an AUC of 0.5.

To account for this potential confounder, we examined the influence of two forms of length information on performance: the number of affected genes and deletion length. We generated two matched test sets with a 1:1 ratio of pathogenic to benign deletions based on the number of affected genes (n=702) and deletion length (n=704), respectively. DosaCNV maintained the best performance on these datasets, achieving AUC values of 0.895 and 0.891, respectively (Supplementary Fig. 2A – 2D; Supplementary Table 1 & 3). Nevertheless, a substantial decrease in performance for all methods was observed when compared to the unmatched test set, suggesting that differences in length distribution contributed to the overly optimistic evaluations in unmatched data.

Furthermore, we noted that controlling for one form of length information may not necessarily account for the other (Supplementary Fig. 1A & B), as pathogenic and benign deletions with similar nucleotide spans might not overlap with a comparable number of genes, and vice versa. To further minimize the impact of length information, we generated a test set matched for both deletion length and the number of affected genes (n=618). This strategy resulted in AUC values of approximately 0.5 for both the length features (deletion length and gene number) in the matched test set and across different length groups (Fig. 2C & D; Supplementary Table 1 & 2), indicating that these two length features could no longer be used for prediction. Even though the overall performance of almost all methods was further reduced, DosaCNV remained the top-performing method (Fig. 2C). DosaCNV outperformed other methods in all length groups, except for the small group (Fig. 2D), where XCNV’s performance was marginally better but not statistically significant (DeLong test p-value = 0.582). By comparing the AUC values of this matched test set to the unmatched test set, we estimated each method’s degree of dependency on length information. CADD-SV demonstrated the greatest dependency with a 0.227 decrease in AUC (unmatched: 0.888, matched: 0.661), while TADA exhibited the smallest dependency with a 0.080 decrease in AUC (unmatched: 0.923, matched: 0.843). DosaCNV had the second smallest dependency, with a 0.091 decrease in AUC (unmatched: 0.968, matched: 0.877).

### DosaCNV enhanced prioritization of putative pathogenic deletions in neurodevelopmental disorder case-control studies

To further assess DosaCNV’s capacity to discriminate between pathogenic and benign deletions, we compared its performance to alternative methods in prioritizing potential pathogenic deletions from case-control studies of neurodevelopmental disorders (NDDs). While ground truth labels for deletions in case-control datasets are unknown, prior research indicates that CNVs may contribute to 15-25% of NDD cases^48,49^. As a result, an effective pathogenic CNV classifier should prioritize these pathogenic CNVs, placing them within the top predictions among all CNVs from cases and controls, with the top-ranked CNVs predominantly depleted in controls. Thus, we assessed the performance of each method based on the enrichment of case-associated deletions within their top predictions, using this as a measure of their ability to prioritize putative pathogenic deletions. Specifically, for the analysis of each dataset, we established the number of top predictions for evaluation as 5% of the total case count (referred to as the top 5% predictions). Based on the conservative assumption that coding CNVs contribute to 10% of cases, with half of these CNVs being deletions.

We examined 16,048 coding deletions from Coe et al., 2014 (dbVar accession number: nstd100)^50^, comprising 29,085 intellectual disability, developmental delay, and autism spectrum disorder (ID/DD/ASD) cases, alongside 19,584 controls. Subsequently, we evaluated 4,230 coding deletions from Zarrei et al., 2019 (dbVar accession number: nstd173)^51^, which included 2,691 NDD cases and 1,769 controls. Within the nstd100 dataset, DosaCNV had the highest enrichment of case-associated deletions among its top 5% predictions (odds ratio = 102.3), which significantly outperforms the second-best method HIS-LOD (odds ratio = 31.1; z-test for log odds ratios p-value = 1.026 x 10^-10^) (Fig. 3A; Supplementary Table 4). Specifically, DosaCNV prioritized 84 more case-associated deletions among top predictions compared to HIS-LOD, suggesting a potential to detect 5.8% (84/1,454) more putative pathogenic deletions. For the nstd173 dataset, DosaCNV again had the highest enrichment of case-associated deletions among its top 5% predictions (odds ratio = 2.12) (Fig. 3B; Supplementary Table 4). However, this did not significantly differ from the next best method TADA (odds ratio = 1.79; z-test for log odds ratios p-value = 0.562), likely attributable to the smaller sample size.

**Figure 3.**
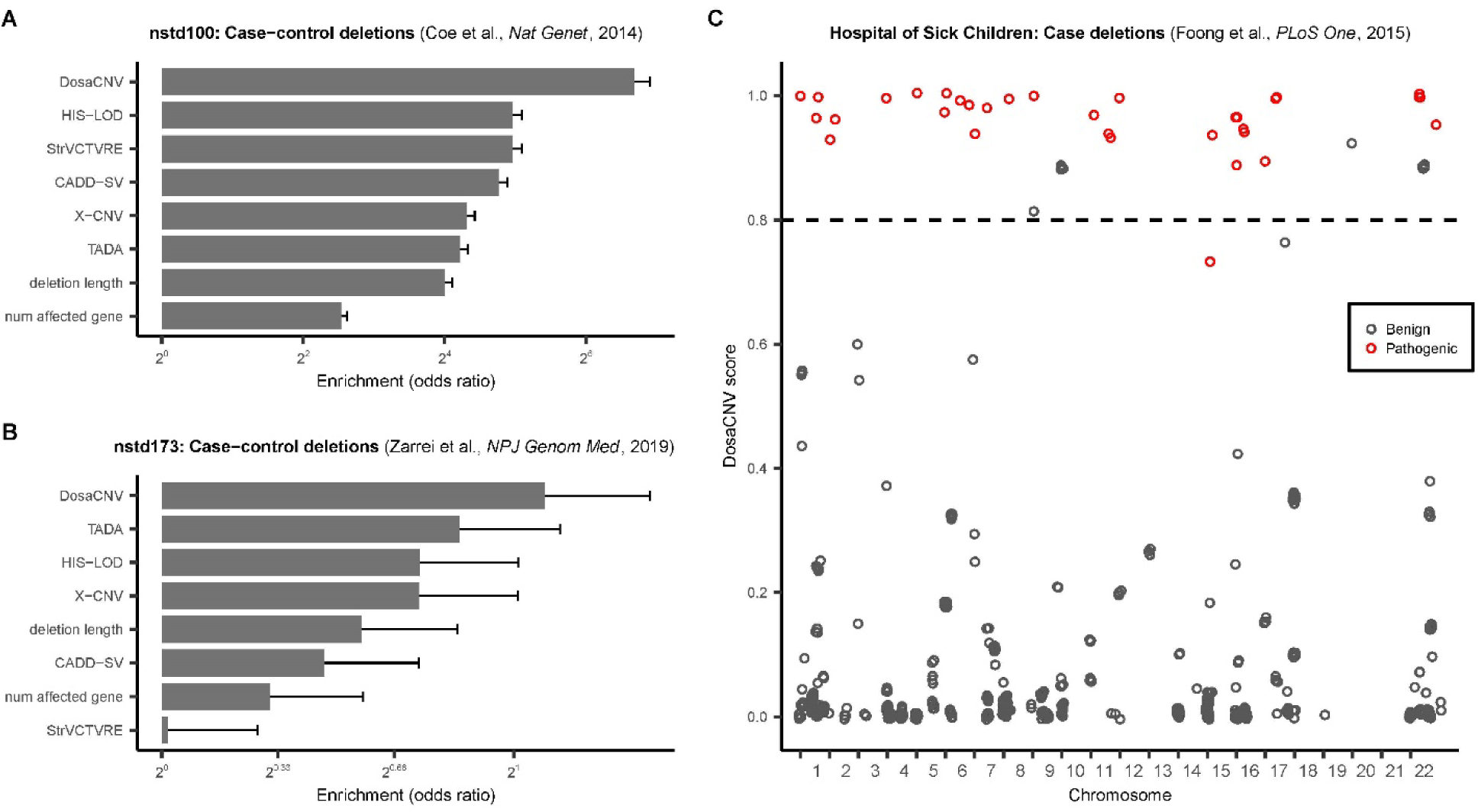
Comparative assessment of DosaCNV and alternative methods for prioritizing potential pathogenic deletions in case-control and curated case-only datasets. (A)(B) Evaluation of prioritization performance in case-control nstd100 (16,048 deletions; 29,085 ID/DD/ASD cases and 19,584 population controls) and nstd173 (4,230 deletions; 2,691 NDD cases and 1,769 family member controls) datasets. The enrichment of case-associated deletions within the top 5% predictions (1,454 for nstd100 and 134 for nstd173) for each method was used as the performance metric. Error bars represent one standard error. (C) DosaCNV scores for 740 clinically curated deletions in a case-only dataset from Foong et al., 2015. A DosaCNV score threshold of 0.8 (indicated by the horizontal dashed line) yields a true positive rate of 0.970 and a false positive rate of 0.028.

Additionally, we investigated the overlap between the two case-control datasets and the ClinVar test set to provide further evidence for DosaCNV’s capacity to prioritize pathogenic deletions. We posited that deletions with over 95% overlap should exhibit similar functional consequences and, therefore, comparable pathogenicity. We identified deletions from the case-control datasets with over 95% overlap with true pathogenic deletions from the ClinVar test set. Then, we assessed whether they were primarily found in cases and prioritized as top predictions by DosaCNV. In the nstd100 dataset, we uncovered 164 overlapping deletions, with 154 observed in cases, substantiating the depletion of most likely pathogenic deletions within controls. Furthermore, 148 of these 164 deletions were included in DosaCNV’s nstd100 top predictions, demonstrating its robust prioritization capability. For the nstd173 dataset, we observed 13 deletions overlapping with true pathogenic deletions, with 11 of them present in cases. Only 4 of the 13 overlapping deletions were among the top 5% predictions of DosaCNV, possibly due to the smaller sample size of the overlapping deletions. Notably, DosaCNV did not prioritize any deletions that overlapped with benign ClinVar deletions as top predictions in either dataset.

Finally, we assessed DosaCNV’s capacity to distinguish pathogenic deletions from benign deletions in patients, a prevalent and challenging clinical scenario when identifying causal variants. We obtained 740 clinically curated coding deletions within 140 DD cases from Foong et al., 2015^52^, which included 33 pathogenic deletions. DosaCNV effectively prioritized pathogenic deletions among benign deletions (Fig. 3C), attaining an AUC of 0.997, which was the highest among all methods (Supplementary Fig. 3). However, the difference in AUC was not statistically significant compared to the second-best method StrVCTVRE (AUC = 0.981; DeLong test p-value = 0.115). With a DosaCNV score threshold of 0.8, the true positive rate reached 0.970, and the false positive rate was 0.028, indicating the accurate classification of almost all pathogenic deletions with only a small fraction of benign deletions misclassified. The precision of DosaCNV was also high (0.604) at this threshold. These results suggest that DosaCNV can accurately distinguish pathogenic deletions from benign deletions in patients, and 0.8 is a reasonable threshold in the practice of clinical variant interpretation.

### DosaCNV-HI outperforms alternative methods in predicting HI genes from multiple sources

To investigate whether the superior performance of DosaCNV at the deletion level is due to more accurate gene haploinsufficiency estimation, we evaluated our model’s ability to distinguish HI and likely HI genes from haplosufficient (HS) genes. First, we assembled six positive sets of HI and likely HI genes, including: 1) 340 ClinGen “gold standard” HI genes^53^, 2) 397 human orthologs of mouse heterozygous knockout lethal (MHKO) genes^54^, 3) 214 high-confidence SFARI ASD genes (SFARI gene score 1)^55^, 4) 432 DDG2P genes sensitive to heterozygous LOF mutations^56^, 5) 72 eyeG2P genes sensitive to heterozygous LOF mutations^56^, and 6) 82 skinG2P genes sensitive to heterozygous LOF mutations^56^. We defined all other genes as the HS gene set. To balance the number of genes and mitigate the impact of CDS length on model performance, we used MatchIt^57^ to match HI and HS genes by CDS length, retaining only matched genes for downstream performance analysis.

To circumvent circularity, we retrained DosaCNV by excluding DDG2P and EDS scores from gene-level input features, as a subset of DDG2P genes was used for gene-level prediction assessment, and the EDS score was trained on a portion of our assembled HI genes. After training, the DosaCNV gene-level model, DosaCNV-HI (Methods), generated HI probability predictions for 20,268 protein-coding genes.

We assessed DosaCNV-HI’s performance against 12 alternative methods across six datasets, encompassing two metrics for LOF mutation intolerance (DeepLOF^29^, and LOEUF^28^), one for missense mutation intolerance (Mis. OEUF^28^), two for both LOF and missense mutation intolerance (RVIS^26^, VIRLoF^58^), and seven for haploinsufficiency (HIS^18^, IS^23^, GHIS^19^, HIPred^20^, Episcore^21^, EDS^22^, pHaplo^24^). The comparison included genes with predictions available from all methods. To evaluate performance robustly within imbalanced sets, we computed the AUC for each method by sampling an equal number of HS genes of matching CDS length as the HI or likely HI gene sets.

DosaCNV-HI significantly surpassed alternative methods in predicting ClinGen HI genes, achieving an AUC of 0.901, which is substantially higher than the second-best method, Episcore, with an AUC of 0.854 (Delong test p-value = 0.019) (Fig. 4A; Supplementary Table 5). DosaCNV-HI consistently achieved high rankings in the remaining five sets of likely HI genes. Notably, even in cases where DosaCNV-HI was not the top performer (ranking second or third), the differences in the AUCs were not statistically significant compared to those of the best methods (Fig. 4B-F; Supplementary Table 5). It is worth noting that the performance of DosaCNV-HI could still be overly optimistic when analyzing all available genes without considering overlap between HI/HS genes and deletions in the ClinVar training set. To address this, we conducted a separate analysis excluding genes overlapped by deletions in the ClinVar training set, and DosaCNV-HI maintained its high-ranking performance across all six gene sets (Supplementary Fig. 4; Supplementary Table 6).

**Figure 4.**
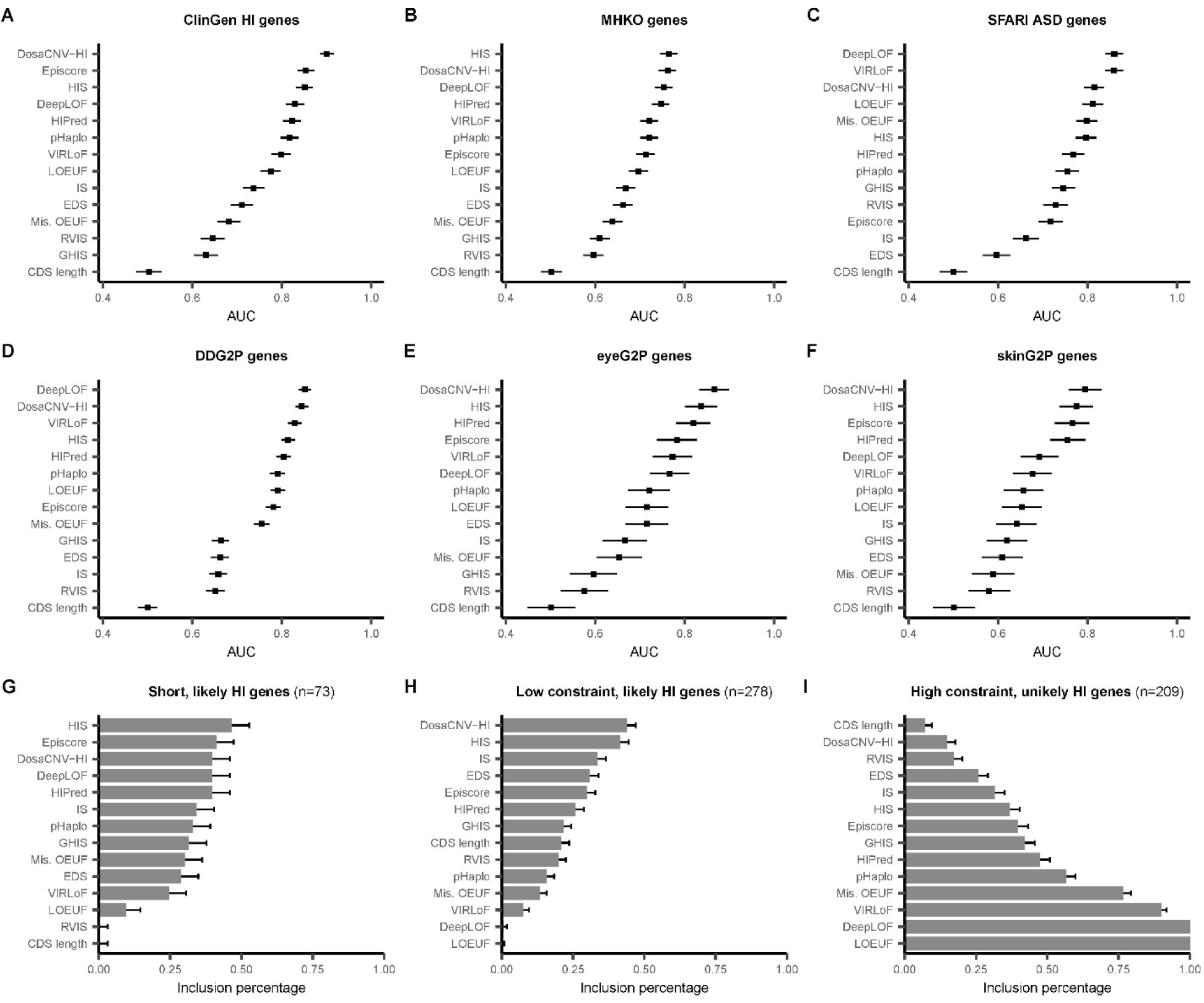
Predictive power of DosaCNV-HI and 12 alternative methods in discriminating HI and likely HI genes from HS genes. AUC values for DosaCNV-HI and alternatives are shown in predicting (A) ClinGen HI genes (n=210/340; 210 out of 340 genes had predictions from all methods), (B) human orthologs of mouse heterozygous knock-out lethal genes (n=315/397), (C) high confidence SFARI ASD genes (n=176/214), (D) DDG2P genes sensitive to heterozygous LOF mutations (n=366/432), (E) eyeG2P genes sensitive to heterozygous LOF mutations (n=59/72), (F) skinG2P genes sensitive to heterozygous LOF mutations (n=76/82). Horizontal black lines denote ± one standard error. Inclusion percentages for (G) “short, likely HI”, (H) “low constraint, likely HI”, and (I) “high constraint, unlikely HI” gene sets within the top 2,402 genes ranked by each method; a higher percentage for “likely HI” sets or a lower percentage for “unlikely HI” sets suggests superior performance. Error bars depict one standard error.

### Evaluating the robustness of DosaCNV-HI and alternative methods in identifying challenging HI and HS genes

Metrics based on evolutionary constraints, such as LOEUF and DeepLOF, demonstrated reasonable performance, indicating that many HI genes are indeed subject to strong purifying selection in human populations. However, these metrics, along with supervised HI gene scores dependent on them, may face challenges in identifying HI genes in three particular cases: 1) short HI genes with limited statistical power for constraint inference^27,28^, 2) HI genes under low evolutionary constraint, including those involved in late-onset disorders (e.g., *BRCA2*, *ATM*, and *ELN*)^33–36^, and 3) falsely predicted HS genes under high evolutionary constraint, such as pleiotropic genes with minor effects on single phenotypes but significant cumulative fitness impacts^31,32^. To evaluate the robustness of our method and alternative approaches in these scenarios, we generated three challenging gene sets.

First, we combined all six HI and likely HI gene sets to form a unified set of 988 genes, aiming to enhance statistical power. From this combined set, we focused on the 758 genes that have predictions across all methods for further analysis. Next, we filtered this set for genes with fewer than 10 expected LOF mutations in human populations^29^, creating a “short, likely HI” gene set (n=73). Second, using an established LOEUF cutoff (<0.35 for constrained genes) and a similar cutoff for DeepLOF (>0.835 for constrained genes), we generated a “low constraint, likely HI” gene set (n=278). Finally, we applied the same cutoffs to identify a set of highly evolutionarily constrained HS genes not implicated in diseases across G2P, Human Phenotype Ontology^59^, ClinVar^42^, and OMIM databases^60^, yielding a “high constraint, unlikely HI” gene set (n=209).

We identified the 2,402 top-ranked genes for each method as predicted HI genes, comparable to the LOEUF cutoff of 0.35, and evaluated performance based on the percentage of each challenging gene set included within the top-ranked genes. An ideal HI gene classifier should maximize the inclusion of genes from the two likely HI gene sets while minimizing the inclusion of genes from the unlikely HI gene set. Among all evaluated methods for the “short, likely HI” gene set, HIS exhibited the highest inclusion percentage (0.466, 34/73), followed by Episcore (0.411, 30/73), and then our method, DosaCNV-HI (0.397, 29/73) (Fig. 4G; Supplementary Table 7). Notably, almost all methods showed a higher inclusion percentage compared to LOEUF. This suggests that these methods, potentially due to the integration of diverse complementary gene-level features, are less affected by the limited statistical power in detecting short, constrained genes. For the “low constraint, likely HI” gene set, DosaCNV-HI and HIS achieved the highest (0.439, 122/278) and second highest (0.414, 115/278) inclusion percentages, respectively (Fig. 4H; Supplementary Table 7). This outcome may be attributed to the fact that these two methods directly model haploinsufficiency instead of evolutionary constraints.

Remarkably, using gene CDS length alone as a predictor resulted in the lowest inclusion percentage (0.072, 15/209) for the “high constraint, unlikely HI” gene set (Fig. 4I; Supplementary Table 7). In contrast, DosaCNV-HI exhibited the second lowest inclusion percentage (0.148, 31/209), with CDS length excluding 7.6% more unlikely HI genes (Fig. 4I). This observation emphasizes a positive correlation between gene haploinsufficiency and CDS length for evolutionarily constrained genes, aligning with prior research that suggests longer genes tend to be more crucial during early developmental stages^61^, with haploinsufficiency potentially serving as an inherent property of these genes^62^. In summary, while DosaCNV-HI may not always be the top-performing method, it consistently ranks highly for predicting challenging HI genes and difficult HS genes.

### Elucidating key features and their contributions to gene haploinsufficiency prediction

We applied Deep SHAP^63^ (SHapley Additive exPlanations) alongside DosaCNV-HI to identify key features determining a gene’s probability of being HI. Based on coalition game theory, Deep SHAP fairly and uniquely allocates an input’s prediction to its features (SHAP values) by computing a weighted average of marginal contributions across all feature combinations. We obtained SHAP values for each feature across 988 HI and likely HI genes using Deep SHAP and sampled 1,000 HS genes as a background set. We then visualized these values using a default SHAP summary plot^63^, offering a global explanation for the entire HI and likely HI gene set (Fig. 5A). The global feature importance is derived from the mean absolute SHAP values over all HI and likely HI genes. Pathogenic SNV presence emerged as the most important feature, as genes sensitive to disrupted SNVs may also be sensitive to dosage changes. Features linked to evolutionary constraints and conservation, such as PLI, LOEUF, and PhastCons, also ranked highly, emphasizing the importance of evolutionary constraints and conservation as common HI gene characteristics (Fig. 5A).

**Figure 5.**
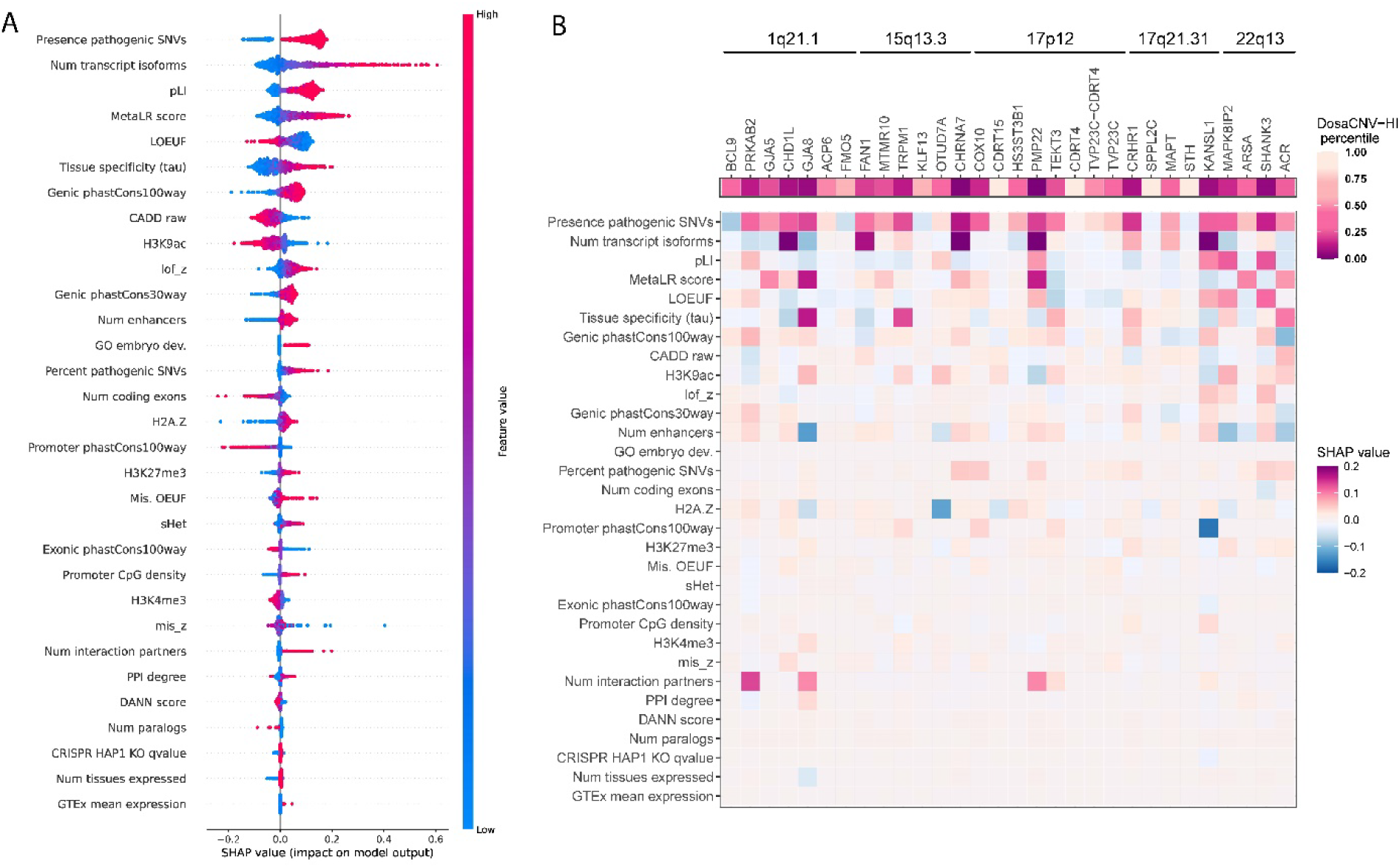
Elucidating gene-level feature contributions to haploinsufficiency predictions via SHAP values. (A) The SHAP summary plot is presented with gene-level features on the y-axis, arranged by global importance (mean absolute SHAP values) in descending order. Each dot represents a specific feature value for a gene within the HI or likely HI gene set (n=988), and the color denotes the actual feature value (red: high; blue: low). The x-axis displays SHAP values, which indicate the impact of each feature on the deviation between a gene’s DosaCNV-HI prediction and the average DosaCNV-HI prediction for background HS genes (n=1000). Positive SHAP values enhance the DosaCNV-HI prediction (genes more likely to be HI), whereas negative values diminish it (genes less likely to be HI). The sum of SHAP values across all features for each gene equals the difference between its predicted HI probability and the expected HI probability of a designated background gene set (HS genes in our case). (B) DosaCNV-HI percentiles are shown for genes overlapping five pathogenic microdeletions, along with their corresponding SHAP values. A gene with a lower DosaCNV-HI percentile is more likely to be HI.

Notably, we observed a positive correlation between the second most important feature, the number of transcript isoforms, and the DosaCNV-HI prediction (Fig. 5A). This finding is in line with a previous study showing that genes with a greater number of isoforms are more likely to be essential genes^64^. The sixth most important feature, tissue specificity (tau), also exhibited a positive correlation with the DosaCNV-HI prediction (Fig. 5A), which implies that HI genes may have greater tissue specificity in expression than HS genes^18^. Epigenetically, higher DosaCNV-HI predictions were associated with stronger signals for repressive histone modifications (H3K27me3) and weaker signals for active modifications (H3K4me3 and H3K9ac) (Fig. 5A). Considering that the majority of epigenetic data were obtained from adult tissues/organs and most HI genes are developmental genes^15^, this result suggests that many HI genes may be epigenetically silenced outside early developmental stages. Intriguingly, we observed a positive correlation between the H2A.Z signal and DosaCNV-HI predictions, although this histone variant is known to facilitate both gene activation and repression^65^. Additionally, several features, such as a greater number of enhancers, increased promoter CpG density, more extensive protein-protein interactions, elevated mean expression levels, and fewer paralogs, contributed to higher DosaCNV-HI predictions (Fig. 5A), aligning with previous studies^18,21,66^.

Deep SHAP also offers local explanations by attributing the HI prediction to individual feature contributions for a specific gene. To demonstrate this, we retrieved coordinates of five recurrent pathogenic microdeletions from ClinGen^67^, and obtained DosaCNV-HI predictions and corresponding SHAP values for 30 protein-coding genes overlapping with these microdeletions (Fig. 5B). DosaCNV-HI identified candidate disease genes with plausible haploinsufficiency mechanisms, consistent with previous findings, such as *CHD1L* and *GJA8* for the 1q21.1 deletion^68,69^, *CHRNA7* for the 15q13.3 deletion^70,71^, *PMP22* for the 17p12 deletion^72^, *CRHR1* and *KANSL1* for the 17q21.31 deletion^73,74^, and *SHANK3* for the 22q13 deletion^75^ (Fig. 5B). Comparing the SHAP values of all features for these candidate genes allows us to understand why they are predicted to have a higher chance of being HI. For example, in the well-documented HI gene *PMP22*, encoding the essential peripheral myelin protein 22 for myelin formation and stability^72^, key contributors to its HI prediction revealed by SHAP values include pathogenic SNVs presence, numerous transcript isoforms, evolutionary constraints, enhancer abundance, and extensive protein interactions (Fig. 5B). These features highlight *PMP22*’s regulatory complexity within the myelin gene network, which implicates the necessity of precise dosage control for its role in proper neural development and function^72^. Similarly, *KANSL1*, another established HI gene involved in chromatin modification and gene regulation^74^, demonstrated a comparable pattern of feature importance, though adapted to its unique biological functions (Fig. 5B). These insights elucidate the rationale behind DosaCNV’s predictions for each gene and deletion, ultimately improving our capacity to explore the biological mechanisms underpinning gene haploinsufficiency and deletion pathogenicity.

## Discussion

In this study, we present DosaCNV, a novel deep MIL model that jointly predicts coding deletion pathogenicity and gene haploinsufficiency. Prior research has examined these predictions independently, without effectively connecting the two. Our approach, for the first time, bridges this gap in a biologically coherent manner, which significantly enhances predictions at both the gene and deletion levels. Moreover, this method enables the evaluation of HI genes’ contribution to deletion pathogenicity and expands the spectrum of learned HI gene features. Additionally, we employ rigorous training and data preprocessing strategies, such as size matching and leave-one-chromosome-out, to strengthen model generalizability and mitigate biases arising from information leakage^76^.

While DosaCNV demonstrated superior performance to other prediction tools in ClinVar data, the results also revealed a notable challenge shared by all methods, including DosaCNV: the dependence of predictions on deletion length. While a strong positive association exists between deletion length and pathogenicity, it is not the primary determinant of pathogenicity. Overreliance on this non-causal association may lead to biased predictions for long benign deletions and short pathogenic deletions^43^. Furthermore, as machine-learning models can easily capture this strong correlation and maintain high accuracy within unmatched datasets, the impact of other features directly involved in the underlying mechanisms of pathogenic deletions might be overshadowed. This emphasizes the importance of taking the length information into account during both model development and performance evaluation.

A robust pathogenic CNV classifier must be generalizable to effectively prioritize putative pathogenic deletions across various disease contexts, addressing the growing accumulation of unlabeled CNV data. DosaCNV, trained on a limited set of pan-disorder deletions, exhibited superior performance in prioritizing putative pathogenic deletions in ID/DD/ASD cases, as evidenced by the highest enrichment of case-associated deletions in predicted pathogenic deletions and its favorable sensitivity in prioritizing clinically curated pathogenic deletions in a real clinical scenario. This generalizability supports DosaCNV’s underlying hypothesis that the pathogenicity of coding deletions predominantly results from affected HI genes, which may share cross-disease commonalities captured by the gene-level features used in the model. However, the performance may also be influenced by the enrichment of ID/DD/ASD patients in our pan-disorder training data. While HI genes often share certain characteristics, their effects may also differ in a disease-dependent manner based on their involvement in distinct biological processes. To account for the variation of gene effects across diseases, DosaCNV is designed for adaptability, enabling easy training with diverse datasets and gene-level features tailored to the specific disease context under investigation.

Data scarcity in predicting gene haploinsufficiency poses a significant challenge for supervised machine-learning methods, with fewer than 1,000 high-confidence HI and likely HI genes available out of an estimated ∼2,900^27,28^. DosaCNV adopts an innovative model design, learning predictive features for haploinsufficiency from affected genes within curated deletions, without requiring gene-level labels. This approach enables a more comprehensive capture of the HI gene landscape, allowing DosaCNV-HI to outperform alternative methods in six HI and likely HI gene sets. However, our model remains dependent on a limited number of clinically curated deletions, which may be subject to ascertainment bias, such as the enrichment of certain types of genes and deletions^45^. Consequently, developing larger and less biased datasets of curated CNVs is crucial for improving novel HI gene discovery and pathogenic deletion prediction.

Evolutionary constraint metrics, such as pLI and LOEUF, have effectively prioritized disease-associated genes based on the idea that deleterious mutations in disease genes are often depleted through purifying selection. While these methods primarily focus on fitness effects, they can also suggest haploinsufficiency, a phenotypic effect, especially for HI genes involved in early developmental processes where carrying a null allele can result in reduced fitness. However, exceptions exist where reduced fitness does not align with haploinsufficiency. For example, mutations in late-onset HI genes may not significantly reduce fitness, and mutations in pleiotropic genes may lead to reduced fitness without causing discernible phenotypic changes^32^. Moreover, evolutionary constraint metrics based on population genetics data are known to be underpowered for short genes with sparse polymorphisms^27–29^. To address these challenges, incorporating gene features orthogonal to evolutionary constraint metrics is crucial. By integrating over 30 gene-level features from diverse biological processes, DosaCNV-HI has demonstrated promising performance in scenarios that pose challenges for traditional evolutionary constraint metrics (Fig. 4G, H, I), underscoring the importance of considering a broader array of gene features to enhance the accuracy and reliability of haploinsufficiency predictions.

To address the limited interpretability of deep learning models, we integrated DosaCNV-HI with a model-specific explanation tool, Deep SHAP^63^, to elucidate key gene features that highlighted the biological characteristics of HI genes (Fig. 5A). Several metrics for evolutionary constraints, such as pLI and LOEUF, emerged as top features, reinforcing the idea that HI gene disruption frequently leads to developmental disorders and is thus subject to strong purifying selection^15,62^. Epigenetically, higher DosaCNV-HI predictions correlated with weaker signals for active histone modification marks (H3K4me3, H3K9ac) and stronger signals for repressive marks (H3K27me3). This supports the hypothesis that a substantial number of HI genes function as developmental transcription factors, which are critical during specific developmental stages and become epigenetically silenced outside these periods^77,78^. Additionally, our results revealed a positive correlation between DosaCNV-HI predictions and H2A.Z signal, which can be attributed to H2A.Z’s prevalent occurrence in bivalent domains within promoters of developmental genes^65^, where HI genes are notably enriched. Regarding regulatory elements, higher enhancer numbers and promoter CpG densities were associated with increased DosaCNV-HI predictions. These characteristics have been associated with essential developmental genes^21,66,79,80^, enabling intricate spatiotemporal expression patterns associated with hypothesized mechanisms of haploinsufficiency^81^.

Certain limitations in our study highlight the need for further refinement to improve the accuracy and comprehensiveness of our method. First, we focused on coding regions in the current study, but the potential contributions of non-coding regions to CNV pathogenicity should not be overlooked^82,83^. Ignoring non-coding regions constrains the interpretation of non-coding CNVs and may impair coding CNV predictions by erroneously assigning pathogenicity to seemingly innocuous coding elements. Second, DosaCNV’s assumption that independent effects of HI genes are the sole contributors to deletion pathogenicity oversimplifies the complex interplay between genomic elements. This simplification fails to account for the epistasis between affected genomic elements and variable penetrance/expressivity arising from other genetic variants, such as SNVs^84–86^. Lastly, we assumed heterozygosity for all deletions in the training and testing sets due to the absence of zygosity information in ClinVar. This assumption is partially supported by the overrepresentation of dominant disease genes and limited overlap with gnomAD benign SVs in ClinVar pathogenic deletions^45^. However, we observed a few pathogenic deletions affecting only single recessive disease genes, suggesting the potential existence of homozygous deletions. Consequently, the accuracy of our HI gene and pathogenic deletion predictions could be impacted, given that DosaCNV was specifically designed to predict haploinsufficiency which is a dominant effect. The future availability of fully phased CNV data will be instrumental in facilitating a more comprehensive understanding of the relationships between the mode of inheritance of disease genes and the pathogenicity of CNVs.

Despite the exclusive focus on deletions in this study, our proposed framework can be suitably adapted to predict the pathogenicity of duplications. This adaptation essentially involves shifting the focus of gene-level prediction from haploinsufficiency to triplosensitivity, achievable by simply utilizing curated duplications as training data. Despite maintaining the same framework structure, a reassessment of predictive features is needed, given the potential discrepancies between HI and TS genes. The integration of appropriate input features will further elucidate the unique characteristics of TS genes.

## Conclusions

In conclusion, DosaCNV is an accurate and robust deep multiple-instance learning framework that simultaneously predicts deletion pathogenicity and gene haploinsufficiency. Traditional methods have often treated these domains in isolation. DosaCNV, however, offers a unified approach to bridge this gap, enhancing predictions at both the deletion and gene levels evidenced in multiple datasets. Furthermore, by integrating the Deep SHAP technique to quantify gene-level feature contributions, DosaCNV not only provides transparent and human-understandable explanations for its predictions but also holds promise in further elucidating the intricate landscape of gene haploinsufficiency. However, challenges persist, notably potential biases in clinically curated CNV datasets, with the ClinVar dataset being an example. Despite our rigorous efforts during model development and evaluation, these biases inevitably limit our model’s precision and overall applicability. As our understanding of previously underexplored genomic regions, such as non-coding regions, expands in tandem with rapid advancements in sequencing technology, we anticipate the emergence of more comprehensive and unbiased CNV catalogs and supporting resources. Such developments will be pivotal in improving the accuracy and reliability of tools like DosaCNV, while also expanding their range of applications.

## Methods

### Details of DosaCNV

We denote **X**_*i*_ as the input for the i-th deletion, which is a matrix of dimensions M x L, where M represents the number of genes and L corresponds to the number of gene-level features. Given that the number of genes varies for each deletion, each **X**_*i*_ is automatically padded to accommodate 100 genes if M < 100 by adding dummy genes with all feature values set to zero. Consequently, **X**_*i*_ assumes a shape of 100 x L. In the fully connected-layer, we aim to model HI probabilities for each of the M genes within the i-th deletion, utilizing their respective gene-level feature vectors (rows of **X**_*i*_). To achieve this, we employ two hidden layers for feature representation learning from **X**_*i*_. The calculations for these layers can be represented by the following equations:

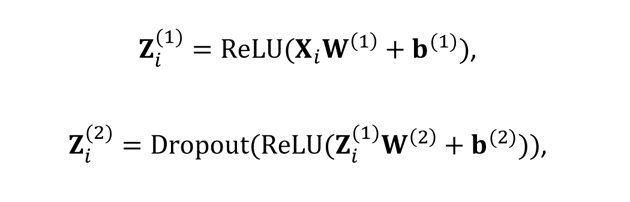

where superscripts (1) and (2) denote the 1st and 2nd hidden layers, respectively. ReLU and Dropout refer to the rectified linear layer^87^ and dropout layer^88^, while W and **b** represent the weight matrix and bias vector, respectively. Z_*j*_ serves as the matrix of hidden units for the i-th deletion, with each row constituting the hidden unit vector for a gene. Subsequently, we incorporate an additional layer to convert Z^(2)^ into HI probabilities (P_HI_) for the genes:

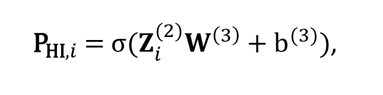

where σ denotes the logistic function, which ensures that the HI probabilities remain within the range of 0 and 1. W(3) and b(3) represent the weight vector and bias term, respectively. P_HI,*j*_ is a vector of length 100, wherein each element serves as the gene-level prediction of haploinsufficiency (DosaCNV-HI score) for a gene or dummy gene within the i-th deletion. To naturally aggregate probabilities within P_HI,*i, j*_ and infer deletion pathogenicity, we introduce a pooling layer that employs the noisy-or function:

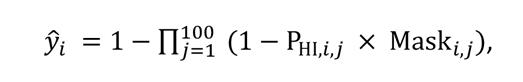

where *Y*_*i*_is the predicted probability of the i-th deletion harboring at least one HI gene and serves as a proxy for deletion pathogenicity. P_HI,*i,j*_ denotes the P_HI_ for the j-th gene or dummy gene within the i-th deletion. Mask_*i,j*_ is a binary indicator, specifying whether the j-th gene is an actual gene (Mask_*i,j*_ = 1) or a dummy gene (Mask_*i,j*_ = 0). This mechanism prevents the propagation of information from dummy genes by setting P_HI,*i,j*_ to zero.

We use the mean binary cross entropy loss to measure the discrepancy between the labels of deletion pathogenicity and predicted pathogenicity probabilities:

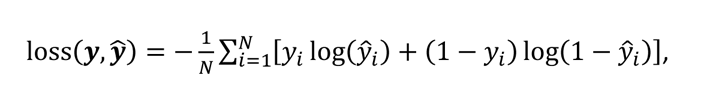

where *y* and *y* represent the vectors of the observed pathogenicity labels and predicted pathogenicity probabilities in a mini-batch of data of size N, respectively.

For model training, we initialize all parameters using He Normal initialization^89^ and employ the Adam algorithm^90^ for mini-batch gradient descent to update model parameters. To mitigate overfitting, we implement early stopping and incorporate an L2 regularization term.

Following the training process, the full model is utilized for variant-level prediction of deletion pathogenicity. Concurrently, all layers of the full model are extracted, excluding the noisy-or pooling layer, to establish a model for gene-level prediction of haploinsufficiency, referred to as DosaCNV-HI.

### ClinVar training, validation, and test set

We obtained the clinically curated rare SVs dataset (GRCh37; dbVar accession number: nstd102) from ClinVar^42^ on May 25, 2021. We filtered for copy number losses/deletions meeting specific criteria: (1) >49 bp in length, (2) not somatic origin, (3) at least 1 bp overlap with a CDS, (4) <101 affected protein-coding genes, and (5) clinical interpretations as pathogenic, likely pathogenic, pathogenic/likely pathogenic, benign, likely benign, or benign/likely benign. Inner start and end coordinates were also required for inclusion. We retrieved coordinates for 20,268 protein-coding genes and their corresponding CDSs from the GENCODE^91^ v38 human basic gene annotation file in GRCh37 and used BEDTools^92^ to determine overlaps with deletions.

To minimize overfitting due to potential ascertainment bias, we identified redundant deletion groups with 95% reciprocal overlap using BEDTools and retained only the shortest deletion within each group. This process yielded 12,002 deletions, which were labeled as pathogenic (1) if they had clinical interpretations as pathogenic, likely pathogenic, or pathogenic/likely pathogenic and benign (0) if they had clinical interpretations as benign, likely benign, or benign/likely benign (6,180 pathogenic and 5,822 benign).

To further minimize overfitting and avoid overly optimistic evaluations due to partial overlaps between deletions in the same chromosome, we adopted a “leave-one-chromosome-out” strategy to partition the deletions into training, validation, and testing sets. The test set included deletions from chromosomes 1, 3, 5, and 7, while the validation set comprised deletions from chromosomes 2, 4, 6, and 8, and the training set contained deletions from the remaining chromosomes. This approach produced 6,678, 2,492, and 2,832 deletions for the training, validation, and testing sets.

We used R package MatchIt^57^ to match pathogenic and benign deletions by the number of affected protein-coding genes in the training and validation sets; only matched pairs were kept. We left the test set unmatched for further analysis of the length impact on performance evaluation. This approach resulted in 2,308 and 1,060 deletions in the final training and validation sets, respectively.

### DosaCNV gene-level features

DosaCNV leverages 33 gene-level features of 20,268 protein-coding genes to predict gene haploinsufficiency and subsequently infer deletion pathogenicity. The gene-level features can be grouped into six main categories: genomics, epigenomics, evolutionary constraints, functional genomics, deleteriousness upon perturbation, and biological networks (Supplementary Data 1). (1) Genomics: We used the Ensembl biomaRt R package^93^ to extract the number of coding exons and transcript isoforms for each gene. We sourced promoter CpG density from a previous study^66^. We retrieved a list of 3,543 human duplicated genes, organized into 945 groups, from the Duplicated Gene Database (DGD)^94^. We determined the number of paralogs for each gene based on the group size of the corresponding gene group and assigned a paralog count of zero for genes absent in the list. (2) Epigenomics: We obtained ChIP-seq broad peaks for H2A.Z, H3K4me3, H3K9ac, and H3K27me3 in promoter regions across cell types, as well as promoter-enhancer interactions predicted by EpiTensor^95^ from a previous study^21^. We calculated the average length of the corresponding ChIP-seq peak in a gene’s promoter region across all cell types to define input features for these epigenomic marks. Additionally, we determined the number of enhancers associated with a gene by averaging the number of predicted promoter-enhancer interactions across cell types. (3) Evolutionary constraint: We extracted lof_z^25^, mis_z^25^, pLI^27^, LOEUF^28^, and Mis. OEUF^28^ from the gnomAD database^28^ v2.1.1. We retrieved SNP-level PhastCons100way vertebrate and PhastCons30way mammalian scores^96^ from the dbNSFP database^97^ v4, and averaged these two metrics across the entire span of each gene to obtain gene-level scores. We sourced promoter and exonic PhastCons100way vertebrate scores from a previous study^66^ and obtained sHet values from the original publication^98^. (4) Functional genomics: We sourced GTEx mean expression^99^, GO terms for embryo development^100^, and tissue specificity (tau)^101^ values from a recent study^29^. We retrieved median gene-level transcripts per million (TPMs) across 54 tissues from the v7 release of the GTEx portal^99^. We counted the number of non-zero expression tissues to determine the number of tissues expressed for each gene. (5) Deleteriousness upon perturbation: We retrieved SNP-level MetaLR^102^, CADD raw^103^, and DANN^104^ scores from the dbNSFP database^97^, and calculated the average scores across all coding exons of each gene. We downloaded a list of 2,545 DDG2P genes from the G2P database^56^ and generated a binary feature to indicate whether a gene is included in this list. We sourced EDS values from the original publication^22^, obtained CRISPR HAP1 cell knockout q-values from a previous study^105^, and downloaded SNV data from the ClinVar^42^. We created a binary feature indicating the presence or absence of pathogenic SNVs in a gene, based on ClinVar SNV data. Additionally, for each gene, we calculated the percentage of pathogenic SNVs relative to the total number of submitted SNVs in ClinVar. (6) Biological networks: We downloaded a dataset of 64,006 binary protein-protein interactions involving 9,094 proteins from the Human Reference Protein Interactome Mapping Project^106^. We counted the number of interaction partners for each gene included in the dataset and assigned a count of zero for genes absent in the dataset. We obtained PPI degree values from a recent study^29^.

We performed mean imputation for missing values and z-standardized all features, excluding binary ones, based on the means and standard deviations derived from genes not overlapping deletions in the ClinVar held-out test set.

### Model training

We mapped 33 gene-level features to each affected gene within deletions, and padded deletions with fewer than 100 genes using dummy genes with all zero feature values within the ClinVar training, validation, and test sets. This process resulted in an input matrix for each deletion with a shape of 100 x 33. We then trained DosaCNV using the ClinVar training set and tuned hyperparameters via grid search based on the ClinVar validation set. The search space included: the number of nodes in each hidden ReLU layer (32, 64, 128, 256); lambda for the L2 penalty (0.006, 0.02, 0.06, 0.2); batch size (16, 32, 64); and Adam learning rate (3×10^-5^, 10^-4^, 3×10^-4^, 10^-3^). The optimal hyperparameters based on the lowest validation loss were 128 and 64 nodes for the first and second hidden ReLU layers, 0.02 for the L2 penalty, a batch size of 32, and a learning rate of 3×10^-4^. Additionally, we set the dropout rate to 0.5^88^.

For gene-level prediction using DosaCNV-HI, we excluded EDS and DDG2P from the gene-level features to avoid potential circularity. DosaCNV was trained on the same ClinVar training set using the same set of optimal hyperparameters from the DosaCNV model for deletion-level prediction.

### Deletion-level evaluation

We obtained deletion-level predictions for comparison from five alternative methods: (1) TADA v1.0.2, available at (https://github.com/jakob-he/TADA), using default settings on the GRCh37 assembly; (2) StrVCTVRE v1.7, accessed from (https://github.com/andrewSharo/StrVCTVRE), employing default settings with all analyzed deletions lifted to GRCh38 using the UCSC liftOver tool; (3) HIS-LOD, in which we retrieved gene-level predictions (HIS) for all genes from the original publication and aggregated them to deletions-level predictions based on the method described therein^18^; (4) X-CNV, downloaded from (https://github.com/kbvstmd/XCNV), with default settings on the GRCh37 assembly; and (5) CADD-SV v1.1 webtool (https://cadd-sv.bihealth.org/score), using default settings on the GRCh37 assembly.

To evaluate performance on the ClinVar held-out test set (n=2,832) and assess the impact of length distribution differences, we compiled a joint set of 2,491 predictions from all five alternative methods, denoting it as the unmatched test set. Using MatchIt, we matched pathogenic and benign deletions in the unmatched test set based on the number of affected protein-coding genes (n=702), deletion length in base pairs (n=704), and both criteria combined (n=618), retaining only matched pairs. To examine performance on deletions of varying lengths, we further divided each of the four sets into four length groups: small (<50 kb), medium (50-100 kb), large (100-1000 kb), and extra-large (>1000 kb). We obtained AUC values and corresponding standard errors for each method on each set using the R package pROC^107^. We visualized the AUC values for all methods using R package ggplot2^108^ and tested the statistical significance for the differences between AUC values using the DeLong test, via the roc.test function in pROC. To estimate each method’s dependency on length information, we subtracted the AUC value of the matched test set from that of the unmatched test set, considering both deletion length and the number of affected genes.

We evaluated the prioritization ability for putative pathogenic deletions by obtaining rare SVs from case-control studies by Coe et al., 2014^50^ (nstd100) with 29,085 ID/DD/ASD cases and 19,584 controls, and Zarrei et al., 2019^51^ (nstd173) with 2,691 NDD cases and 1,769 family member controls. We applied the same filtering conditions with the ClinVar data, excluding the condition for clinical interpretation. The resulting nstd100 and nstd173 sets contained 16,528 and 4,500 coding deletions, respectively, with 16,048 and 4,230 deletions having predictions from all methods. Since these datasets lack pathogenicity labels, we calculated the enrichment of case-associated deletions in each method’s top predictions using odds ratio as the performance measure. For each dataset, we defined the number of top predictions for evaluation as 5% of the total case count (referred to as the top 5% predictions), assuming that coding CNVs contribute to 10% of cases, half of which are deletions. This approach resulted in 1,454 and 134 top predictions for nstd100 and nstd173, respectively. We visualized the odds ratios by ggplot2 and tested the statistical significance for the differences between odds ratios by transforming them to log odds ratios and then using the z-test.

To further assess our method, we determined the inclusion percentage of the nstd100 and nstd173 deletions that had over 95% reciprocal overlap with curated pathogenic deletions from ClinVar, within our method’s top predictions. Specifically, we found 164 nstd100 deletions and 13 nstd173 deletions had over 95% reciprocal overlap with pathogenic deletions in the ClinVar held-out test set.

To evaluate performance in a clinical scenario, we acquired clinically curated rare SVs from 140 developmental delay cases reported by Foong et al., 2015^52^. After filtering, 740 coding deletions, including 33 pathogenic ones, remained. We generated predictions for these deletions and visualized them in a scatter plot using ggplot2. Additionally, 663 of the 740 deletions had predictions from all methods. Based on the joint set of 663 deletions, we determined the AUC values for our method and the alternative methods using pROC.

### Gene-level evaluation

To evaluate the performance of DosaCNV-HI and 12 alternative methods— including DeepLOF^29^, LOEUF^28^, Mis.OEUF^28^, RVIS^26^, VIRLoF^58^, HIS^18^, IS^23^, GHIS^19^, HIPred^20^, Episcore^21^, EDS^22^, and pHaplo^24^—in predicting HI and likely HI genes, we obtained LOEUF and Mis.OEUF from the gnomAD database v2.1.1 and gene scores for all other methods from their respective original publications.

We retrieved six HI and likely HI gene sets for evaluation: (1) 340 “gold standard” HI genes with sufficient evidence for haploinsufficiency (ClinGen HI score 3) from the ClinGen gene dosage sensitivity map^53^; (2) 397 human orthologs of mouse heterozygous knockout lethal genes from the GitHub repository for gnomAD (https://github.com/macarthur-lab/gnomAD_lof/); (3) 214 high-confidence ASD genes (SFARI gene score 1) from the SFARI GENE database^55^; (4) 432 DDG2P genes with the monoallelic autosomal allelic requirement and absent gene product mutation consequence from the G2P database^56^; (5) 72 eyeG2P genes with the same requirements as DDG2P genes from the G2P database; and (6) 82 skinG2P genes with the same requirements as DDG2P genes from the G2P database. We conducted analyses using genes with predictions from all methods, resulting in 210, 315, 176, 360, 59, and 76 HI/likely HI genes remaining within the six gene lists, respectively. All other protein-coding genes were considered as the HS gene set.

To mitigate the imbalance between the six HI or likely HI gene sets and the HS gene set, and to control the influence of gene length on performance evaluation, we adopted a matching approach. For each HI or likely HI gene set, we used MatchIt to pair each HI gene with an HS gene of a similar CDS length to obtain a matched gene set. Subsequently, we utilized pROC to compute the AUC values and corresponding standard errors for each method within the matched gene sets. We visualized the AUC values for all methods using ggplot2 and tested the statistical significance for the differences between AUC values using the DeLong test.

For the evaluation of challenging HI and HS genes, we aggregated the six gene sets into a single HI and likely HI gene list (n=988). From genes with predictions across all methods (n=758), we generated a “short, likely HI” gene set (n=73) by filtering genes with fewer than 10 expected LOF mutations from the gnomAD database. Using established LOEUF (<0.35 for constrained genes)^28^ and DeepLOF (>0.835 for constrained genes)^29^ cutoffs for constrained genes, we created a “low constraint, likely HI” gene set (n=278). Applying the same cutoffs, we generated a “high constraint, unlikely HI” gene set (n=209) for constrained HS genes not implicated in diseases across the G2P, Human Phenotype Ontology^59^, ClinVar, and OMIM databases^60^. Subsequently, we defined the top 2,402 genes predicted by each method as the top predictions for each method (based on cutoff: LOEUF <0.35), representing the predicted HI genes. We calculated the inclusion percentages with corresponding standard errors of the three challenging gene sets in each method’s top predictions as the performance measure and visualized these inclusion percentages using ggplot2.

### Feature importance

To elucidate the key features driving DosaCNV-HI gene-level predictions, we employed the DeepExplainer function from the SHAP v0.41.0 Python library^63^. We generated a background set, consisting of 31 gene-level features for 1,000 randomly selected HS genes, resulting in a 1,000 x 31 matrix where each row represents a gene-level feature vector for a specific gene. The explainer was initialized with the DosaCNV-HI model and the defined background set, allowing the calculation of conditional expectations for each gene-level feature and the corresponding conditional expectation of HI predictions for the 1,000 randomly sampled HS genes. Subsequently, we generated an input set containing 31 gene-level features for 988 HI and likely HI genes, which we fed into the explainer to obtain the SHAP values. The output was a 988 x 31 SHAP value matrix, where each row represents the SHAP values for corresponding gene-level feature values of a particular gene. We used the default bee swarm summary plot from the package to visualize the global importance and impact distributions of gene-level features. The features on the y-axis were ranked in descending order of global importance, which the package calculated as the mean absolute value for the corresponding column in the output SHAP value matrix.

We employed the same approach to generate SHAP values for 30 protein-coding genes overlapping pathogenic microdeletions at 1q21.1 (distal, BP3-BP4), 15q13.3 (BP4-BP5), 17p12 (HNPP), 17q21.31, and 22q13, with coordinates retrieved from the ClinGen database in the GRCh37 assembly. We visualized the SHAP values and percentiles of DosaCNV-HI predictions for these affected genes as a heatmap using ggplot2.

## Availability of data and materials

The DosaCNV program and the precomputed DosaCNV-HI gene scores can be accessed at https://github.com/Zhihan-Leo-Liu/DosaCNV. Datasets used in this study, along with the source code for analysis can be found at https://github.com/Zhihan-Leo-Liu/DosaCNV/analysis/. Additionally, the sources and download links for all gene-level features are provided in Supplementary Data 1.

## Acknowledgments

The authors thank Mingfu Shao and Xinru Zhang for helpful discussions.

## Funding

Research reported in this publication was supported by the National Institute of General Medical Sciences of the National Institutes of Health under Award Number R35GM142560 and by startup funds from Pennsylvania State University. The content is solely the responsibility of the author and does not necessarily represent the official views of the National Institutes of Health.

## Author information

**Authors and Affiliations**

Department of Biology, Pennsylvania State University, University Park, PA 16802, USA Zhihan Liu & Yi-Fei Huang

Huck Institutes of the Life Sciences, Pennsylvania State University, University Park, PA 16802, USA Zhihan Liu & Yi-Fei Huang

Molecular, Cellular, and Integrative Biosciences Program, Pennsylvania State University, University Park, PA 16802, USA

Zhihan Liu

## Contributions

ZL implemented the DosaCNV program and conducted all analyses. YH conceived and supervised the project. ZL and YH prepared and reviewed the manuscript.

## Corresponding authors

Correspondence to Zhihan Liu or Yi-Fei Huang.

## Ethics declarations

**Ethics approval and consent to participate**

Not applicable

## Consent for publication

Not applicable

## Competing interests

The authors declare no competing interests.

## Additional materials

**Additional file 1**

Additional file 1. docx

Supplementary Figures 1-4 and Supplementary Tables 1-7

## Additional file 2

Additional file 2.xls

The sources and download links for all gene-level features used in this study. Referred to as Supplementary Data 1 in the main text.

## Notes

### Competing Interest Statement

The authors have declared no competing interest.

### Summary of Updates

I have updated a few numerical values in the manuscript after fixing a minor bug in the DosaCNV program.

## References

1. Zhang, F., Gu, W., Hurles, M. E. & Lupski, J. R. Copy Number Variation in Human Health, Disease, and Evolution. Annu. Rev. Genomics Hum. Genet. 10, 451–481 (2009).

2. Conrad, D. F. et al. Origins and functional impact of copy number variation in the human genome. Nature 464, 704–712 (2010).

3. Hastings, P. J., Lupski, J. R., Rosenberg, S. M. & Ira, G. Mechanisms of change in gene copy number. Nat. Rev. Genet. 10, 551–564 (2009).

4. Liu, P., Carvalho, C. M. B., Hastings, P. J. & Lupski, J. R. Mechanisms for recurrent and complex human genomic rearrangements. Curr. Opin. Genet. Dev. 22, 211–220 (2012).

5. Perry, G. H. et al. Diet and the evolution of human amylase gene copy number variation. Nat. Genet. 39, 1256–1260 (2007).

6. Leffler, E. M., et al. Resistance to malaria through structural variation of red blood cell invasion receptors. Science 356, eaam6393 (2017).

7. Malhotra, D. & Sebat, J. CNVs: Harbingers of a Rare Variant Revolution in Psychiatric Genetics. Cell 148, 1223–1241 (2012).

8. McCarroll, S. A. et al. Deletion polymorphism upstream of IRGM associated with altered IRGM expression and Crohn’s disease. Nat. Genet. 40, 1107–1112 (2008).

9. McCarroll, S. A. et al. Donor-recipient mismatch for common gene deletion polymorphisms in graft-versus-host disease. Nat. Genet. 41, 1341–1344 (2009).

10. Cooper, G. M. et al. A copy number variation morbidity map of developmental delay. Nat. Genet. 43, 838–846 (2011).

11. Pinto, D. et al. Functional impact of global rare copy number variation in autism spectrum disorders. Nature 466, 368–372 (2010).

12. McCarthy, S. E. et al. Microduplications of 16p11.2 are associated with schizophrenia. Nat. Genet. 41, 1223–1227 (2009).

13. Shao, X. et al. Copy number variation is highly correlated with differential gene expression: a pan-cancer study. BMC Med. Genet. 20, 175 (2019).

14. Weischenfeldt, J., Symmons, O., Spitz, F. & Korbel, J. O. Phenotypic impact of genomic structural variation: insights from and for human disease. Nat. Rev. Genet. 14, 125–138 (2013).

15. Rice, A. M. & McLysaght, A. Dosage sensitivity is a major determinant of human copy number variant pathogenicity. Nat. Commun. 8, 14366 (2017).

16. Seidman, J. G. & Seidman, C. Transcription factor haploinsufficiency: when half a loaf is not enough. J. Clin. Invest. 109, 451–455 (2002).

17. Veitia, R. A. & Potier, M. C. Gene dosage imbalances: action, reaction, and models. Trends Biochem. Sci. 40, 309–317 (2015).

18. Huang, N., Lee, I., Marcotte, E. M. & Hurles, M. E. Characterising and Predicting Haploinsufficiency in the Human Genome. PLOS Genet. 6, e1001154 (2010).

19. Steinberg, J., Honti, F., Meader, S. & Webber, C. Haploinsufficiency predictions without study bias. Nucleic Acids Res. 43, e101 (2015).

20. Shihab, H. A., Rogers, M. F., Campbell, C. & Gaunt, T. R. HIPred: an integrative approach to predicting haploinsufficient genes. Bioinformatics 33, 1751–1757 (2017).

21. Han, X. et al. Distinct epigenomic patterns are associated with haploinsufficiency and predict risk genes of developmental disorders. Nat. Commun. 9, 2138 (2018).

22. Wang, X. & Goldstein, D. B. Enhancer Domains Predict Gene Pathogenicity and Inform Gene Discovery in Complex Disease. Am. J. Hum. Genet. 106, 215–233 (2020).

23. Khurana, E., Fu, Y., Chen, J. & Gerstein, M. Interpretation of Genomic Variants Using a Unified Biological Network Approach. PLOS Comput. Biol. 9, e1002886 (2013).

24. Collins, R. L. et al. A cross-disorder dosage sensitivity map of the human genome. Cell 185, 3041–3055.e25 (2022).

25. Samocha, K. E. et al. A framework for the interpretation of de novo mutation in human disease. Nat. Genet. 46, 944–950 (2014).

26. Petrovski, S., Wang, Q., Heinzen, E. L., Allen, A. S. & Goldstein, D. B. Genic Intolerance to Functional Variation and the Interpretation of Personal Genomes. PLoS Genet. 9, e1003709 (2013).

27. Lek, M. et al. Analysis of protein-coding genetic variation in 60,706 humans. Nature 536, 285–291 (2016).

28. Karczewski, K. J. et al. The mutational constraint spectrum quantified from variation in 141,456 humans. Nature 581, 434–443 (2020).

29. LaPolice, T. M. & Huang, Y.-F. *A deep learning framework for predicting human essential genes from population and functional genomic data*. (2021) doi:10.1101/2021.12.21.473690.

30. Ruderfer, D. M. et al. Patterns of genic intolerance of rare copy number variation in 59,898 human exomes. Nat. Genet. 48, 1107–1111 (2016).

31. Simons, Y. B., Bullaughey, K., Hudson, R. R. & Sella, G. A population genetic interpretation of GWAS findings for human quantitative traits. PLOS Biol. 16, e2002985 (2018).

32. Fuller, Z. L., Berg, J. J., Mostafavi, H., Sella, G. & Przeworski, M. Measuring intolerance to mutation in human genetics. Nat. Genet. 51, 772–776 (2019).

33. Wooster, R. et al. Identification of the breast cancer susceptibility gene BRCA2. Nature 378, 789–792 (1995).

34. Roy, R., Chun, J. & Powell, S. N. BRCA1 and BRCA2: different roles in a common pathway of genome protection. Nat. Rev. Cancer 12, 68–78 (2012).

35. Raybould, M. C., Birley, A. J. & Hultén, M. Molecular variation of the human elastin (ELN) gene in a normal human population. Ann. Hum. Genet. 59, 149–161 (1995).

36. Lu, S. et al. Atm-haploinsufficiency enhances susceptibility to carcinogen-induced mammary tumors. Carcinogenesis 27, 848–855 (2006).

37. Ganel, L., Abel, H. J., FinMetSeq Consortium & Hall, I. M. SVScore: an impact prediction tool for structural variation. Bioinformatics 33, 1083–1085 (2017).

38. Geoffroy, V., et al. AnnotSV: an integrated tool for structural variations annotation. Bioinformatics 34, 3572–3574 (2018).

39. Riggs, E. R. et al. Technical standards for the interpretation and reporting of constitutional copy number variants: a joint consensus recommendation of the American College of Medical Genetics and Genomics (ACMG) and the Clinical Genome Resource (ClinGen). Genet. Med. Off. J. Am. Coll. Med. Genet. 22, 245–257 (2020).

40. Amores, J. Multiple instance classification: Review, taxonomy and comparative study. Artif. Intell. 201, 81–105 (2013).

41. Pearl, J. Probabilistic Reasoning in Intelligent Systems: Networks of Plausible Inference. (Morgan Kaufmann, 1988).

42. Landrum, M. J. et al. ClinVar: improving access to variant interpretations and supporting evidence. Nucleic Acids Res. 46, D1062–D1067 (2018).

43. Skelly, A. C., Dettori, J. R. & Brodt, E. D. Assessing bias: the importance of considering confounding. Evid.-Based Spine-Care J. 3, 9–12 (2012).

44. Hertzberg, J., Mundlos, S., Vingron, M. & Gallone, G. TADA—a machine learning tool for functional annotation-based prioritisation of pathogenic CNVs. Genome Biol. 23, 67 (2022).

45. Sharo, A. G., Hu, Z., Sunyaev, S. R. & Brenner, S. E. StrVCTVRE: A supervised learning method to predict the pathogenicity of human genome structural variants. Am. J. Hum. Genet. 109, 195–209 (2022).

46. Zhang, L. et al. X-CNV: genome-wide prediction of the pathogenicity of copy number variations. Genome Med. 13, 132 (2021).

47. Kleinert, P. & Kircher, M. A framework to score the effects of structural variants in health and disease. Genome Res. gr.275995.121 (2022) doi:10.1101/gr.275995.121.

48. Girirajan, S. & Eichler, E. E. Phenotypic variability and genetic susceptibility to genomic disorders. Hum. Mol. Genet. 19, R176–R187 (2010).

49. Miller, D. T. et al. Consensus Statement: Chromosomal Microarray Is a First-Tier Clinical Diagnostic Test for Individuals with Developmental Disabilities or Congenital Anomalies. Am. J. Hum. Genet. 86, 749–764 (2010).

50. Coe, B. P. et al. Refining analyses of copy number variation identifies specific genes associated with developmental delay. Nat. Genet. 46, 1063–1071 (2014).

51. Zarrei, M. et al. A large data resource of genomic copy number variation across neurodevelopmental disorders. NPJ Genomic Med. 4, 26 (2019).

52. Foong, J., Girdea, M., Stavropoulos, J. & Brudno, M. Prioritizing Clinically Relevant Copy Number Variation from Genetic Interactions and Gene Function Data. PLOS ONE 10, e0139656 (2015).

53. Riggs, E. R. et al. Copy number variant discrepancy resolution using the ClinGen dosage sensitivity map results in updated clinical interpretations in ClinVar. Hum. Mutat. 39, 1650–1659 (2018).

54. Bult, C. J., Blake, J. A., Smith, C. L., Kadin, J. A. & Richardson, J. E. Mouse Genome Database (MGD) 2019. Nucleic Acids Res. 47, D801–D806 (2019).

55. Abrahams, B. S. et al. SFARI Gene 2.0: a community-driven knowledgebase for the autism spectrum disorders (ASDs). Mol. Autism 4, 36 (2013).

56. Thormann, A. et al. Flexible and scalable diagnostic filtering of genomic variants using G2P with Ensembl VEP. Nat. Commun. 10, 2373 (2019).

57. Ho, D., Imai, K., King, G. & Stuart, E. A. MatchIt: Nonparametric Preprocessing for Parametric Causal Inference. J. Stat. Softw. 42, 1–28 (2011).

58. Abramovs, N., Brass, A. & Tassabehji, M. GeVIR is a continuous gene-level metric that uses variant distribution patterns to prioritize disease candidate genes. Nat. Genet. 52, 35–39 (2020).

59. Köhler, S. et al. The Human Phenotype Ontology in 2021. Nucleic Acids Res. 49, D1207–D1217 (2020).

60. Amberger, J., Bocchini, C. A., Scott, A. F. & Hamosh, A. McKusick’s Online Mendelian Inheritance in Man (OMIM®). Nucleic Acids Res. 37, D793–D796 (2009).

61. Lopes, I., Altab, G., Raina, P. & de Magalhães, J. P. Gene Size Matters: An Analysis of Gene Length in the Human Genome. Front. Genet. 12, (2021).

62. Zug, R. Developmental disorders caused by haploinsufficiency of transcriptional regulators: a perspective based on cell fate determination. Biol. Open 11, bio058896 (2022).

63. Lundberg, S. & Lee, S.-I. A Unified Approach to Interpreting Model Predictions. Preprint at 10.48550/arXiv.1705.07874 (2017).

64. Yong Ryu, J., Uk Kim, H. & Yup Lee, S. Human genes with a greater number of transcript variants tend to show biological features of housekeeping and essential genes. Mol. Biosyst. 11, 2798–2807 (2015).

65. Ku, M. et al. H2A.Z landscapes and dual modifications in pluripotent and multipotent stem cells underlie complex genome regulatory functions. Genome Biol. 13, R85 (2012).

66. Boukas, L., Bjornsson, H. T. & Hansen, K. D. Promoter CpG Density Predicts Downstream Gene Loss-of-Function Intolerance. Am. J. Hum. Genet. 107, 487–498 (2020).

67. Rehm, H. L. et al. ClinGen — The Clinical Genome Resource. N. Engl. J. Med. 372, 2235–2242 (2015).

68. Harvard, C. et al. Understanding the impact of 1q21.1 copy number variant. Orphanet J. Rare Dis. 6, 54 (2011).

69. Prokudin, I. et al. Exome sequencing in developmental eye disease leads to identification of causal variants in GJA8, CRYGC, PAX6 and CYP1B1. *Eur. J. Hum. Genet.* **22**, 907–915 (2014).

70. Hoppman-Chaney, N., Wain, K., Seger, P., Superneau, D. & Hodge, J. Identification of single gene deletions at 15q13.3: further evidence that CHRNA7 causes the 15q13.3 microdeletion syndrome phenotype. Clin. Genet. 83, 345–351 (2013).

71. Gillentine, M. & Schaaf, C. P. The Human Clinical Phenotypes of Altered CHRNA7 Copy Number. Biochem. Pharmacol. 97, 352–362 (2015).

72. Li, J., Parker, B., Martyn, C., Natarajan, C. & Guo, J. The PMP22 Gene and Its Related Diseases. Mol. Neurobiol. 47, 673–698 (2013).

73. Wray, C. D. 17q21.31 Microdeletion associated with infantile spasms. Eur. J. Med. Genet. 56, 59–61 (2013).

74. Koolen, D. A. et al. Mutations in the chromatin modifier gene KANSL1 cause the 17q21.31 microdeletion syndrome. Nat. Genet. 44, 639–641 (2012).

75. Bozdagi, O. et al. Haploinsufficiency of the autism-associated Shank3 gene leads to deficits in synaptic function, social interaction, and social communication. Mol. Autism 1, 15 (2010).

76. Whalen, S., Schreiber, J., Noble, W. S. & Pollard, K. S. Navigating the pitfalls of applying machine learning in genomics. Nat. Rev. Genet. 23, 169–181 (2022).

77. Creyghton, M. P. et al. H2AZ Is Enriched at Polycomb Complex Target Genes in ES Cells and Is Necessary for Lineage Commitment. Cell 135, 649–661 (2008).

78. Yang, X. et al. Silencing of developmental genes by H3K27me3 and DNA methylation reflects the discrepant plasticity of embryonic and extraembryonic lineages. Cell Res. 28, 593–596 (2018).

79. Frankel, N. et al. Phenotypic robustness conferred by apparently redundant transcriptional enhancers. Nature 466, 490–493 (2010).

80. Norris, M., Lovell, S. & Delneri, D. Characterization and Prediction of Haploinsufficiency Using Systems-Level Gene Properties in Yeast. G3 GenesGenomesGenetics 3, 1965–1977 (2013).

81. Johnson, A. F., Nguyen, H. T. & Veitia, R. A. Causes and effects of haploinsufficiency. Biol. Rev. 94, 1774–1785 (2019).

82. Spielmann, M., Lupiáñez, D. G. & Mundlos, S. Structural variation in the 3D genome. Nat. Rev. Genet. 19, 453–467 (2018).

83. Bhartiya, D. & Scaria, V. Genomic variations in non-coding RNAs: Structure, function and regulation. Genomics 107, 59–68 (2016).

84. Carvalho, C. M. B. et al. Dosage Changes of a Segment at 17p13.1 Lead to Intellectual Disability and Microcephaly as a Result of Complex Genetic Interaction of Multiple Genes. Am. J. Hum. Genet. 95, 565–578 (2014).

85. Girirajan, S. et al. Phenotypic Heterogeneity of Genomic Disorders and Rare Copy-Number Variants. N. Engl. J. Med. 367, 1321–1331 (2012).

86. Davies, R. W. et al. Using common genetic variation to examine phenotypic expression and risk prediction in 22q11.2 deletion syndrome. Nat. Med. 26, 1912–1918 (2020).

87. Nair, V. & Hinton, G. E. Rectified Linear Units Improve Restricted Boltzmann Machines.

88. Srivastava, N., Hinton, G., Krizhevsky, A., Sutskever, I. & Salakhutdinov, R. Dropout: A Simple Way to Prevent Neural Networks from Overfitting.

89. He, K., Zhang, X., Ren, S. & Sun, J. Delving Deep into Rectifiers: Surpassing Human-Level Performance on ImageNet Classification. Preprint at htp://arxiv.org/abs/1502.01852 (2015).

90. Kingma, D. P. & Ba, J. Adam: A Method for Stochastic Optimization. Preprint at htp://arxiv.org/abs/1412.6980 (2017).

91. Frankish, A. et al. GENCODE reference annotation for the human and mouse genomes. Nucleic Acids Res. 47, D766–D773 (2019).

92. Quinlan, A. R. & Hall, I. M. BEDTools: a flexible suite of utilities for comparing genomic features. Bioinformatics 26, 841–842 (2010).

93. Durinck, S., Spellman, P. T., Birney, E. & Huber, W. Mapping identifiers for the integration of genomic datasets with the R/Bioconductor package biomaRt. Nat. Protoc. 4, 1184–1191 (2009).

94. Ouedraogo, M. et al. The Duplicated Genes Database: Identification and Functional Annotation of Co-Localised Duplicated Genes across Genomes. PLoS ONE 7, e50653 (2012).

95. Zhu, Y. et al. Constructing 3D interaction maps from 1D epigenomes. Nat. Commun. 7, 10812 (2016).

96. Siepel, A. et al. Evolutionarily conserved elements in vertebrate, insect, worm, and yeast genomes. Genome Res. 15, 1034–1050 (2005).

97. Liu, X., Li, C., Mou, C., Dong, Y. & Tu, Y. dbNSFP v4: a comprehensive database of transcript-specific functional predictions and annotations for human nonsynonymous and splice-site SNVs. Genome Med. 12, 103 (2020).

98. Cassa, C. A. et al. Estimating the Selective Effects of Heterozygous Protein Truncating Variants from Human Exome Data. Nat. Genet. 49, 806–810 (2017).

99. The GTEx Consortium atlas of genetic regulatory effects across human tissues. Science 369, 1318– 1330 (2020).

100. The Gene Ontology Resource: 20 years and still GOing strong. Nucleic Acids Res. 47, D330–D338 (2019).

101. Yanai, I. et al. Genome-wide midrange transcription profiles reveal expression level relationships in human tissue specification. Bioinformatics 21, 650–659 (2005).

102. Dong, C. et al. Comparison and integration of deleteriousness prediction methods for nonsynonymous SNVs in whole exome sequencing studies. Hum. Mol. Genet. 24, 2125–2137 (2015).

103. Kircher, M. et al. A general framework for estimating the relative pathogenicity of human genetic variants. Nat. Genet. 46, 310–315 (2014).

104. Quang, D., Chen, Y. & Xie, X. DANN: a deep learning approach for annotating the pathogenicity of genetic variants. Bioinformatics 31, 761–763 (2015).

105. Blomen, V. A. et al. Gene essentiality and synthetic lethality in haploid human cells. Science 350, 1092–1096 (2015).

106. Luck, K. et al. A reference map of the human binary protein interactome. Nature 580, 402–408 (2020).

107. Robin, X. et al. pROC: an open-source package for R and S+ to analyze and compare ROC curves. BMC Bioinformatics 12, 77 (2011).

108. Wickham, H. ggplot2: Elegant Graphics for Data Analysis. (Springer, 2009). doi:10.1007/978-0-387-98141-3.

