## Supplementary Figures 1-4 and Supplementary Tables 1-7 for "Deep multiple-instance learning accurately predicts gene haploinsufficiency and deletion pathogenicity"

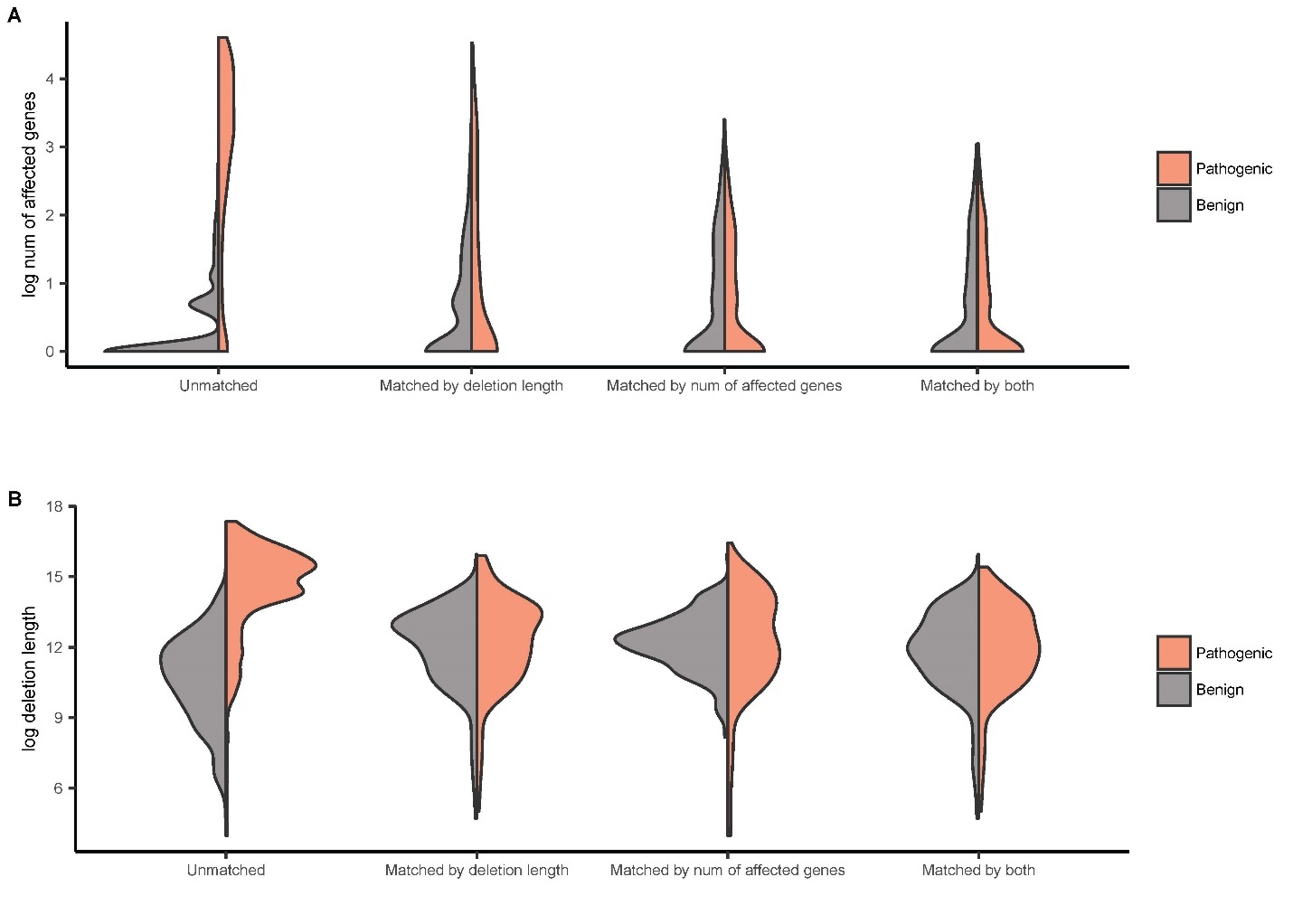

**Supplementary Figure 1.** Length distributions of pathogenic and benign deletions in the ClinVar held-out test set under four matching conditions between pathogenic and benign deletions: unmatched (no matching), matched by deletion length (in base pairs), matched by number of affected genes, and matched by both criteria. (A) Representation based on the natural log-transformed number of affected genes. (B) Representation using the natural log-transformed deletion length.

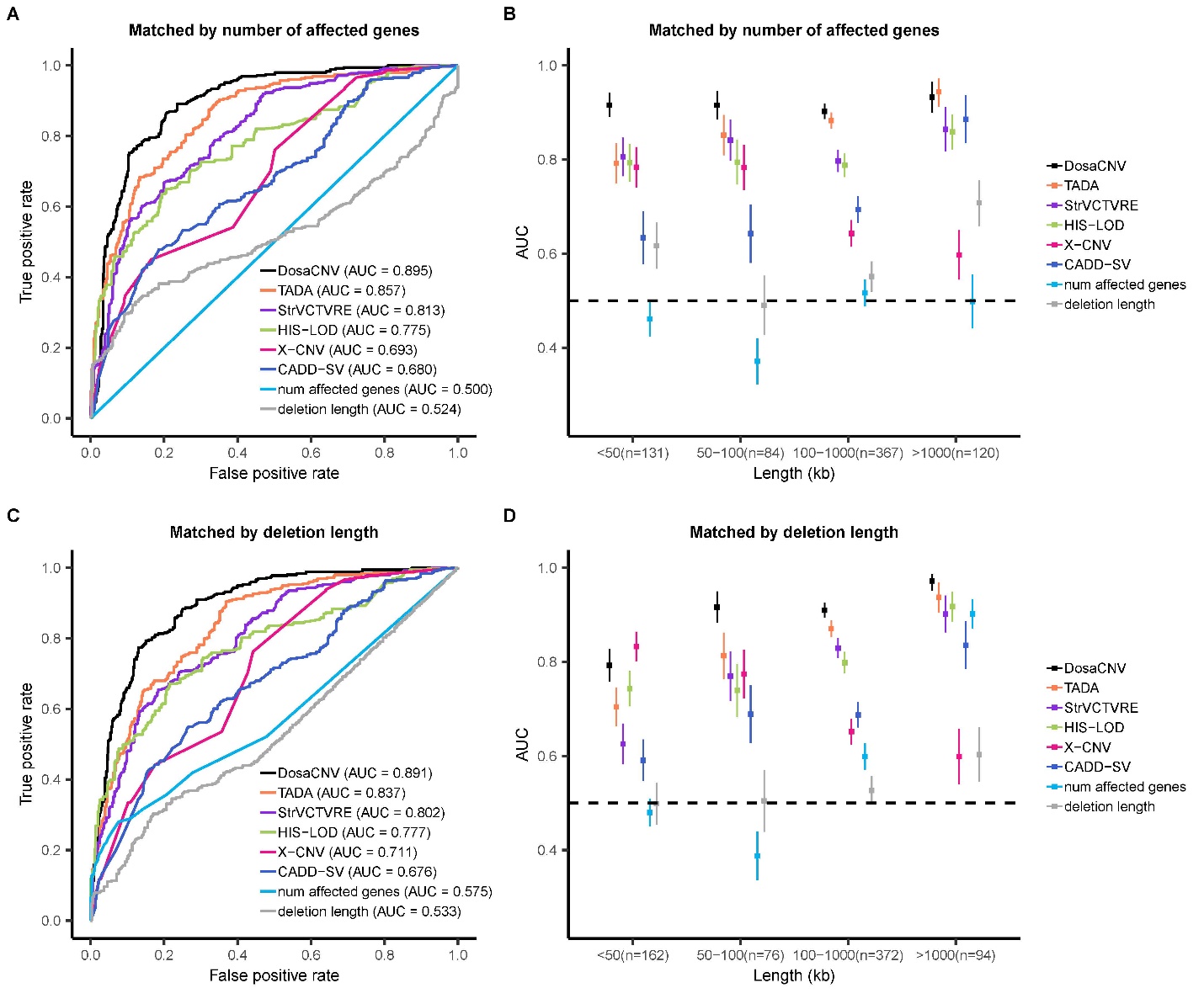

**Supplementary Figure 2.** Performance comparison of DosaCNV and alternative methods in predicting the pathogenicity of deletions from the ClinVar held-out test set. This analysis is based on two matched subsets from the ClinVar test set: matched by the number of affected genes and matched by deletion length. For each set, comparisons are presented for the entire set and four deletion subgroups based on length: small (<50 kb), medium (50-100 kb), large (100-1000 kb), and extra-large (>1000 kb). (A)(B) Test set matched by the number of affected genes (n=702). (C)(D) Test set matched by deletion length (n=704). Vertical lines denote ± one standard error. Horizontal dashed lines indicate an AUC of 0.5.

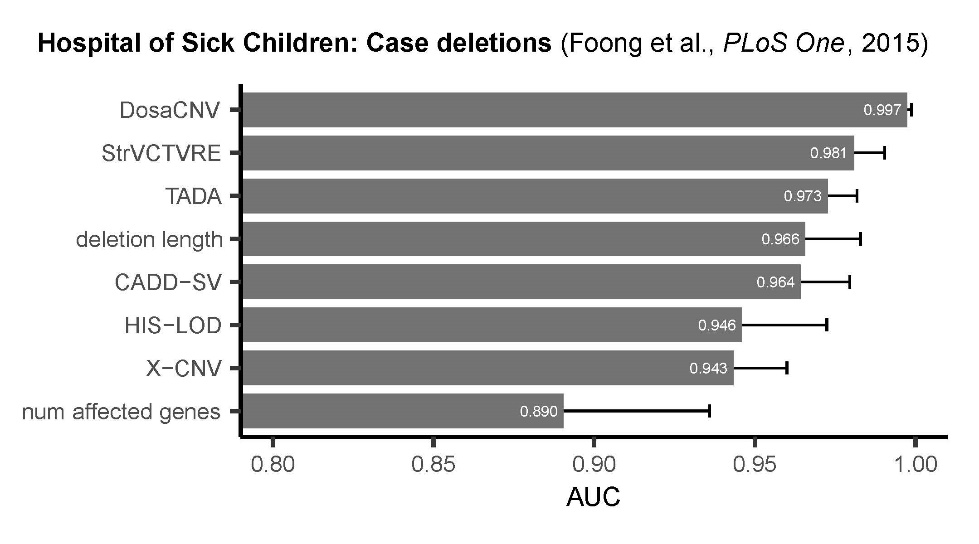

**Supplementary Figure 3**. Performance comparison of DosaCNV and alternative methods in predicting the pathogenicity of 740 clinically curated deletions from 140 DD cases, sourced from a case-only study according to Foong et al., 2015. Of the 740 deletions, 663 had predictions from all methods. AUC comparisons between methods were conducted using this joint set of 663 deletions. Error bars represent one standard error.

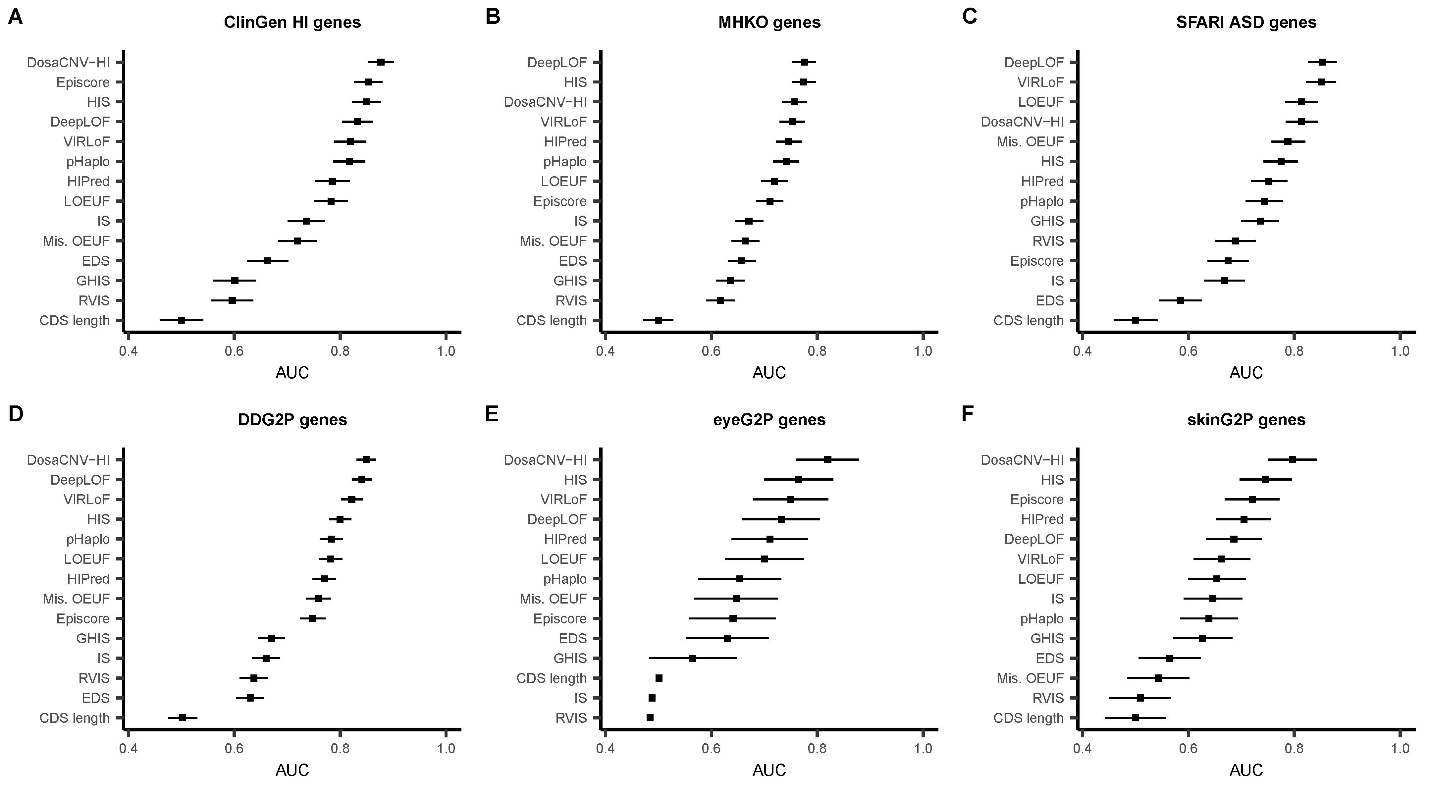

**Supplementary Figure 4.** Predictive power of DosaCNV-HI and 12 alternative methods in discriminating HI and likely HI genes from HS genes. All genes overlapped by deletions in the ClinVar training set were excluded from this analysis. AUC values for DosaCNV-HI and alternatives are shown in predicting (A) ClinGen HI genes (n=99/340; 99 out of 340 genes had predictions from all methods), (B) human orthologs of mouse heterozygous knock-out lethal genes (n=209/397), (C) high confidence SFARI ASD genes (n=98/214), (D) DDG2P genes sensitive to heterozygous LOF mutations (n=219/432), (E) eyeG2P genes sensitive to heterozygous LOF mutations (n=26/72), and (F) skinG2P genes sensitive to heterozygous LOF mutations (n=51/82). Horizontal black lines denote ± one standard error.

**Supplementary Table 1.** AUC values (± standard errors) of DosaCNV and alternative methods in predicting the pathogenicity of deletions from the ClinVar held-out test set, under four matching conditions between pathogenic and benign deletions: unmatched, matched by the number of affected genes, matched by deletion length, and matched by both criteria. Highlighted are the AUC values of the top two methods for each condition, with DeLong test p-values indicating the statistical significance of differences.

|  | Unmatched | Matched by number of affected genes | Matched by deletion length | Matched by both |
| --- | --- | --- | --- | --- |
| DosaCNV | **0.968±0.003** | **0.895±0.012** | **0.891±0.012** | **0.877±0.014** |
| TADA | 0.923±0.005 | **0.857±0.014** | **0.837±0.015** | **0.843±0.016** |
| StrVCTVRE | 0.915±0.006 | 0.813±0.016 | 0.802±0.016 | 0.764±0.019 |
| HIS-LOD | **0.939±0.005** | 0.775±0.017 | 0.777±0.017 | 0.752±0.019 |
| X-CNV | 0.838±0.008 | 0.693±0.019 | 0.711±0.019 | 0.688±0.021 |
| CADD-SV | 0.887±0.007 | 0.680±0.020 | 0.676±0.020 | 0.661±0.022 |
| num affected genes | 0.886±0.007 | 0.500±0.020 | 0.575±0.020 | 0.500±0.021 |
| deletion length | 0.912±0.006 | 0.524±0.023 | 0.533±0.022 | 0.520±0.023 |
| p-value | **6.398x10^-15^** | **1.729x10^-3^** | **3.746x10^-5^** | **1.171x10^-2^** |

**Supplementary Table 2.** AUC values (± standard errors) of DosaCNV and alternative methods in predicting the pathogenicity of deletions from the ClinVar held-out test set, segmented into four deletion length groups: small (<50 kb), medium (50-100 kb), large (100-1000 kb), and extra-large (>1000 kb). Comparisons are made under two matching conditions between pathogenic and benign deletions: unmatched, matched by both the number of affected genes and deletion length. Highlighted are the AUC values of the top two methods per group and condition, with associated DeLong test p-values for statistical significance of differences.

|  | Matching condition | Small  (<50 kb) | Medium  (50-100 kb) | Large  (100-1000 kb) | Extra-large  (>1000 kb) |
| --- | --- | --- | --- | --- | --- |
| DosaCNV | Unmatched | **0.817±0.021** | **0.901±0.028** | **0.922±0.011** | **0.984±0.009** |
|  | Both | **0.809±0.035** | **0.897±0.039** | **0.909±0.018** | **0.930±0.033** |
| TADA | Unmatched | 0.703±0.029 | **0.820±0.042** | **0.881±0.015** | 0.946±0.030 |
|  | Both | 0.708±0.041 | **0.797±0.052** | **0.891±0.018** | **0.928±0.037** |
| StrVCTVRE | Unmatched | 0.610±0.029 | 0.804±0.042 | 0.836±0.018 | 0.942±0.027 |
|  | Both | 0.621±0.044 | 0.735±0.058 | 0.822±0.024 | 0.826±0.053 |
| HIS-LOD | Unmatched | 0.742±0.030 | 0.743±0.042 | 0.817±0.020 | **0.951±0.017** |
|  | Both | 0.732±0.039 | 0.713±0.059 | 0.783±0.027 | 0.830±0.049 |
| X-CNV | Unmatched | **0.821±0.022** | 0.748±0.040 | 0.664±0.023 | 0.649±0.044 |
|  | Both | **0.832±0.032** | 0.740±0.055 | 0.642±0.031 | 0.493±0.066 |
| CADD-SV | Unmatched | 0.595±0.031 | 0.659±0.049 | 0.722±0.023 | 0.849±0.048 |
|  | Both | 0.587±0.045 | 0.615±0.066 | 0.671±0.031 | 0.848±0.053 |
| num affected genes | Unmatched | 0.514±0.021 | 0.389±0.038 | 0.617±0.026 | 0.946±0.011 |
|  | Both | 0.494±0.029 | 0.488±0.048 | 0.501±0.032 | 0.533±0.066 |
| deletion length | Unmatched | 0.562±0.038 | 0.559±0.053 | 0.661±0.026 | 0.872±0.025 |
|  | Both | 0.505±0.046 | 0.517±0.067 | 0.534±0.033 | 0.592±0.066 |
| p-value | Unmatched | **8.802x10^-1^** | **1.260x10^-2^** | **4.392x10^-3^** | **2.322x10^-2^** |
| p-value | Both | **5.815x10^-1^** | **1.222x10^-2^** | **2.823x10^-1^** | **9.591x10^-1^** |

**Supplementary Table 3.** AUC values (± standard errors) of DosaCNV and alternative methods in predicting the pathogenicity of deletions from the ClinVar held-out test set, segmented into four deletion length groups: small (<50 kb), medium (50-100 kb), large (100-1000 kb), and extra-large (>1000 kb). Comparisons are made under two matching conditions between pathogenic and benign deletions: matched by the number of affected genes, and matched by deletion length. Highlighted are the AUC values of the top two methods per length group and matching condition, with associated DeLong test p-values for statistical significance of differences.

|  | Matching condition | Small  (<50 kb) | Medium  (50-100 kb) | Large  (100-1000 kb) | Extra-large  (>1000 kb) |
| --- | --- | --- | --- | --- | --- |
| DosaCNV | Number of affected genes | **0.916±0.026** | **0.915±0.031** | **0.902±0.016** | **0.933±0.034** |
|  | Deletion length | **0.792±0.036** | **0.916±0.034** | **0.910±0.016** | **0.972±0.017** |
| TADA | Number of affected genes | 0.792±0.043 | **0.852±0.044** | **0.883±0.017** | **0.944±0.030** |
|  | Deletion length | 0.704±0.042 | **0.813±0.050** | **0.870±0.018** | **0.937±0.033** |
| StrVCTVRE | Number of affected genes | **0.806±0.042** | 0.841±0.044 | 0.797±0.024 | 0.864±0.048 |
|  | Deletion length | 0.625±0.044 | 0.769±0.054 | 0.829±0.021 | 0.901±0.040 |
| HIS-LOD | Number of affected genes | 0.793±0.041 | 0.794±0.048 | 0.788±0.025 | 0.859±0.038 |
|  | Deletion length | 0.743±0.038 | 0.739±0.057 | 0.798±0.023 | 0.917±0.032 |
| X-CNV | Number of affected genes | 0.784±0.044 | 0.783±0.048 | 0.643±0.029 | 0.597±0.054 |
|  | Deletion length | **0.833±0.032** | 0.774±0.052 | 0.651±0.028 | 0.599±0.060 |
| CADD-SV | Number of affected genes | 0.634±0.057 | 0.642±0.063 | 0.694±0.029 | 0.886±0.052 |
|  | Deletion length | 0.591±0.045 | 0.689±0.062 | 0.687±0.028 | 0.835±0.052 |
| num affected genes | Number of affected genes | 0.462±0.038 | 0.371±0.049 | 0.517±0.029 | 0.498±0.058 |
|  | Deletion length | 0.480±0.030 | 0.388±0.053 | 0.599±0.029 | 0.902±0.032 |
| deletion length | Number of affected genes | 0.617±0.050 | 0.490±0.064 | 0.551±0.033 | 0.708±0.050 |
|  | Deletion length | 0.499±0.046 | 0.505±0.067 | 0.527±0.031 | 0.603±0.059 |
| p-value | Number of affected genes | **1.848x10^-3^** | **7.395x10^-2^** | **2.445x10^-1^** | **7.397x10^-1^** |
| p-value | Deletion length | **3.196x10^-1^** | **4.271x10^-3^** | **3.212x10^-2^** | **3.308x10^-1^** |

**Supplementary Table 4.** Enrichment of case-associated deletions within the top 5% predictions (1,454 for nstd100 and 134 for nstd173) for each method, represented as odds ratios (OR) and log(OR) (± standard errors). The log(OR) of the top two performing methods for each dataset are highlighted, along with their corresponding z-test p-values for the statistical significance of their differences.

|  | nstd100 | | |  | | nstd173 | | |
| --- | --- | --- | --- | --- | --- | --- | --- | --- |
|  | | OR | log(OR) | |  | | OR | log(OR) |
| DosaCNV | | 102.264 | **4.628±0.158** | |  | | 2.121 | **0.752±0.210** |
| TADA | | 18.578 | 2.922±0.079 | |  | | 1.792 | **0.583±0.201** |
| StrVCTVRE | | 31.102 | **3.437±0.095** | |  | | 1.007 | 0.007±0.182 |
| HIS-LOD | | 31.091 | 3.437±0.095 | |  | | 1.657 | 0.505±0.198 |
| X-CNV | | 19.809 | 2.986±0.081 | |  | | 1.655 | 0.504±0.198 |
| CADD-SV | | 27.098 | 3.299±0.090 | |  | | 1.372 | 0.316±0.190 |
| num affected genes | | 5.808 | 1.759±0.060 | |  | | 1.233 | 0.210±0.187 |
| deletion length | | 16.002 | 2.773±0.076 | |  | | 1.476 | 0.390±0.193 |
| p-value | |  | **1.026x10^-10^** | |  | |  | **5.619x10^-1^** |

**Supplementary Table 5.** AUC values (± standard errors) of DosaCNV-HI and alternative methods in differentiating six positive sets of HI and likely HI genes from HS genes. The sets include: (1) ClinGen HI genes (n=210/340; 210 out of 340 genes had predictions from all methods), (2) human orthologs of mouse heterozygous knock-out lethal genes (n=315/397), (3) high confidence SFARI ASD genes (n=176/214), (4) DDG2P genes sensitive to heterozygous LOF mutations (n=366/432), (5) eyeG2P genes sensitive to heterozygous LOF mutations (n=59/72), and (6) skinG2P genes sensitive to heterozygous LOF mutations (n=76/82). Highlighted are the AUC values of the top two methods for each positive set, with DeLong test p-values indicating the statistical significance of differences. If our method is not among the top two, we highlight the AUC values of our method and the leading method, alongside the corresponding DeLong test results.

|  | ClinGen HI genes | MHKO genes | SFARI ASD genes | DDG2P genes | eyeG2P genes | skinG2P genes |
| --- | --- | --- | --- | --- | --- | --- |
| DosaCNV-HI | **0.901±0.015** | **0.762±0.019** | **0.815±0.022** | **0.845±0.014** | **0.867±0.034** | **0.795±0.036** |
| Episcore | **0.854±0.018** | 0.713±0.021 | 0.717±0.027 | 0.781±0.017 | 0.783±0.045 | 0.766±0.039 |
| HIS | 0.851±0.018 | **0.765±0.019** | 0.797±0.024 | 0.814±0.016 | **0.837±0.036** | **0.775±0.038** |
| DeepLOF | 0.830±0.020 | 0.753±0.019 | **0.860±0.020** | **0.852±0.014** | 0.766±0.044 | 0.692±0.042 |
| HIPred | 0.823±0.020 | 0.747±0.019 | 0.768±0.025 | 0.804±0.016 | 0.820±0.039 | 0.756±0.039 |
| pHaplo | 0.818±0.020 | 0.721±0.020 | 0.755±0.026 | 0.791±0.017 | 0.721±0.047 | 0.657±0.044 |
| VIRLoF | 0.799±0.021 | 0.721±0.020 | 0.859±0.020 | 0.829±0.015 | 0.773±0.044 | 0.677±0.043 |
| LOEUF | 0.776±0.023 | 0.697±0.021 | 0.812±0.023 | 0.791±0.017 | 0.716±0.047 | 0.653±0.044 |
| IS | 0.737±0.024 | 0.668±0.021 | 0.662±0.029 | 0.658±0.020 | 0.666±0.050 | 0.641±0.045 |
| EDS | 0.711±0.025 | 0.662±0.021 | 0.596±0.030 | 0.662±0.020 | 0.716±0.048 | 0.609±0.046 |
| Mis. OEUF | 0.682±0.026 | 0.639±0.022 | 0.798±0.024 | 0.755±0.018 | 0.654±0.051 | 0.589±0.047 |
| RVIS | 0.645±0.027 | 0.596±0.023 | 0.728±0.027 | 0.652±0.020 | 0.576±0.053 | 0.580±0.046 |
| GHIS | 0.630±0.027 | 0.610±0.022 | 0.746±0.026 | 0.664±0.020 | 0.597±0.053 | 0.620±0.045 |
| CDS length | 0.503±0.028 | 0.502±0.023 | 0.500±0.031 | 0.500±0.021 | 0.501±0.054 | 0.501±0.047 |
| p-value | **1.890x10^-2^** | **8.647x10^-1^** | **6.620x10^-2^** | **6.459x10^-1^** | **4.424x10^-1^** | **5.869x10^-1^** |

**Supplementary Table 6.** AUC values (± standard errors) of DosaCNV-HI and alternative methods in differentiating six positive sets of HI and likely HI genes from HS genes, excluding genes in positive sets overlapped by deletions in the ClinVar training set. The sets include: (1) ClinGen HI genes (n=99/340; 99 out of 340 genes had predictions from all methods), (2) human orthologs of mouse heterozygous knock-out lethal genes (n=209/397), (3) high confidence SFARI ASD genes (n=98/214), (4) DDG2P genes sensitive to heterozygous LOF mutations (n=219/432), (5) eyeG2P genes sensitive to heterozygous LOF mutations (n=26/72), and (6) skinG2P genes requiring heterozygous LOF mutations (n=51/82). Highlighted are the AUC values of the top two methods for each positive set, with DeLong test p-values indicating the statistical significance of differences. If our method is not among the top two, we highlight the AUC values of our method and the leading method, alongside the corresponding DeLong test results.

|  | ClinGen HI  genes | MHKO genes | SFARI ASD genes | DDG2P genes | eyeG2P genes | skinG2P genes |
| --- | --- | --- | --- | --- | --- | --- |
| DosaCNV-HI | **0.877±0.025** | **0.757±0.024** | **0.814±0.030** | **0.849±0.018** | **0.820±0.059** | **0.797±0.046** |
| Episcore | **0.853±0.027** | 0.711±0.025 | 0.675±0.039 | 0.748±0.024 | 0.641±0.081 | 0.721±0.051 |
| HIS | 0.850±0.027 | 0.774±0.023 | 0.775±0.033 | 0.800±0.021 | **0.765±0.065** | **0.746±0.049** |
| DeepLOF | 0.833±0.029 | **0.776±0.023** | **0.853±0.027** | **0.841±0.019** | 0.732±0.073 | 0.686±0.052 |
| HIPred | 0.785±0.032 | 0.746±0.024 | 0.752±0.034 | 0.770±0.022 | 0.711±0.072 | 0.705±0.052 |
| pHaplo | 0.817±0.030 | 0.742±0.024 | 0.743±0.035 | 0.784±0.022 | 0.654±0.079 | 0.639±0.055 |
| VIRLoF | 0.819±0.030 | 0.753±0.024 | 0.851±0.028 | 0.822±0.020 | 0.750±0.071 | 0.663±0.054 |
| LOEUF | 0.783±0.033 | 0.719±0.025 | 0.814±0.031 | 0.782±0.022 | 0.700±0.074 | 0.654±0.054 |
| IS | 0.736±0.035 | 0.671±0.026 | 0.669±0.038 | 0.660±0.026 | 0.488±0.083 | 0.646±0.055 |
| EDS | 0.663±0.039 | 0.657±0.026 | 0.585±0.041 | 0.630±0.026 | 0.630±0.079 | 0.564±0.059 |
| Mis. OEUF | 0.719±0.037 | 0.664±0.027 | 0.788±0.032 | 0.759±0.023 | 0.647±0.079 | 0.543±0.059 |
| RVIS | 0.596±0.040 | 0.617±0.027 | 0.689±0.038 | 0.637±0.026 | 0.484±0.082 | 0.509±0.058 |
| GHIS | 0.601±0.040 | 0.636±0.027 | 0.736±0.036 | 0.670±0.026 | 0.565±0.082 | 0.627±0.055 |
| CDS length | 0.500±0.041 | 0.500±0.028 | 0.500±0.041 | 0.502±0.028 | 0.501±0.082 | 0.500±0.058 |
| p-value | **4.261x10^-1^** | **4.193x10^-1^** | **2.278x10^-1^** | **6.791x10^-1^** | **4.393x10^-1^** | **3.321x10^-1^** |

**Supplementary Table 7.** Inclusion percentages (± standard errors) for “short, likely HI” (n=73), “low constraint, likely HI” (n=278), and “high constraint, unlikely HI” gene sets (n=209) within the top 2,402 genes ranked by each method. Higher percentages for “likely HI” sets and a lower percentage for “unlikely HI” set suggest superior performance. Highlighted are percentages of the top two methods for each gene set and their corresponding z-test p-values for the statistical significance of their differences.

|  | Short,  likely HI genes | Low constraint, likely HI genes | High constraint, unlikely HI genes |
| --- | --- | --- | --- |
| DeepLOF | 0.397±0.062 | 0.007±0.011 | 1.000±0.000 |
| DosaCNV-HI | **0.397±0.062** | **0.439±0.031** | **0.148±0.029** |
| HIS | **0.466±0.061** | **0.414±0.031** | 0.368±0.036 |
| Episcore | 0.411±0.062 | 0.299±0.030 | 0.397±0.036 |
| HIPred | 0.397±0.062 | 0.259±0.029 | 0.474±0.036 |
| Mis. OEUF | 0.301±0.061 | 0.133±0.024 | 0.766±0.028 |
| pHaplo | 0.329±0.062 | 0.158±0.025 | 0.565±0.035 |
| EDS | 0.288±0.061 | 0.309±0.030 | 0.258±0.034 |
| IS | 0.342±0.062 | 0.335±0.030 | 0.316±0.035 |
| VIRLoF | 0.247±0.060 | 0.076±0.020 | 0.900±0.018 |
| GHIS | 0.315±0.062 | 0.216±0.028 | 0.421±0.036 |
| LOEUF | 0.096±0.050 | 0.000±0.009 | 1.000±0.000 |
| RVIS | 0.000±0.032 | 0.198±0.027 | 0.172±0.030 |
| CDS length | 0.000±0.032 | 0.209±0.027 | **0.072±0.024** |
| p-value | **4.034x10^-1^** | **5.483x10^-1^** | **1.240x10^-2^** |
